# PredicTF: a tool to predict bacterial transcription factors in complex microbial communities

**DOI:** 10.1101/2021.01.28.428666

**Authors:** Lummy Maria Oliveira Monteiro, Joao Saraiva, Rodolfo Brizola Toscan, Peter F Stadler, Rafael Silva-Rocha, Ulisses Nunes da Rocha

**Affiliations:** Helmholtz Center for Environmental Research (UFZ), Leipzig, Germany; Bioinformatics Group, Institute of Computer Science, Universität Leipzig, Leipzig, Germany; Ribeirão Preto Medical School (FMRP), University of São Paulo (USP), Ribeirão Preto, Brazil

**Keywords:** Gene regulation, Transcription factors, Deep Learning, Transcription factor database, Microbial Communities

## Abstract

Transcription Factors (TFs) are proteins that control the flow of genetic information by regulating cellular gene expression. Here we describe PredicTF, a first platform supporting the prediction and classification of novel bacterial TF in complex microbial communities. We evaluated PredicTF using a two-step approach. First, we tested PredictTF’s ability to predict TFs for the genome of an environmental isolate. In the second evaluation step, PredicTF was used to predict TFs in a metagenome and 11 metatranscriptomes recovered from a community performing anaerobic ammonium oxidation (anammox) in a bioreactor. PredicTF is open source pipeline available at https://github.com/mdsufz/PredicTF.

## Background

The functional potential of microbial communities can be determined by the genetic content of its constituent members. However, genetic content alone does not guarantee that a given function or enzymatic reaction will be performed [1]. In this scenario, Transcription Factor proteins (TFs) play a central and critical role in gene regulation. These proteins are responsible for optimizing proteins and structural RNAs and the subsequent levels of metabolites and other properties, ensuring the survival and adaptation of organisms to the most diverse types of stress and environmental changes [2]. The activity of bacterial TFs is modulated by environmental signals (e.g. changes in the oxygen condition, temperature, pH or the lack of a specific substrate) [3]. Additionally, for many promoters, combinations of transcription factors work together to integrate different signals [2,4]. TFs can also work with other DNA-binding proteins whose primary role is to sculpt the bacterial folded chromosome [2,5]. Knowledge of the TFs profile expressed by an organism is the first step to better understand the regulatory network that controls protein expression in an organism or a community.

Since TFs may determine when and which genes are expressed, profiling TFs can help understand the regulation of gene expression and to build regulatory networks in complex microbial communities. Further, defining which factors control gene expression may offer insights into the mechanisms controlling ecosystem processes and even interactions between species of a microbial community. However, current TF databases are focused on single or small groups of genomes. These databases are largely manually curated based on literature evidence and pairwise sequence comparison of genomes from model organisms. Examples of these databases include RegulonDB for *Escherichia coli* K-12 [6], DBTBS for *Bacillus subtilis* [7], FlyBase for *Drosophila* [8], and FTFD for fungal species [9]. DBD [10], is a database generated from the prediction of TFs from 150 sequenced genomes from across the tree of life. Unfortunately, DBD has not been updated for more than 9 years.

One of the major goals in the manipulation of microbiomes for ecological and biotechnological applications is to control the outcome of their functions [11]. As TFs are key to potentially control which genes are expressed, one of the best ways to study and understand gene regulation in a microbiome may be to profile its TFs. To date, no platform supports prediction and classification of novel bacterial TF from ‘omics data recovered from microbial communities.

Deep Learning approaches have been used to predict DNA sequence affinities [12] and to identify TF-binding sites in humans [13]. Although deep learning has been used in gene regulation, it has never been used to predict bacterial TFs. Further, the need for a user-friendly tool for prediction of TFs that could assist in gene regulation analysis motivated the development of PredicTF. PredicTF is a deep learning tool used to predict and identify TFs from full protein-length sequences. Further, we constructed a robust database for bacterial transcriptional factors (BacTFDB) that was used to train our Deep Learning model.

## Results and Discussion

PredictTF is a command line software for prediction of novel transcription factors from genomic and metagenomic data. We created a bacterial transcription factor database (BacTFDB) by merging and manually curating TFs present in CollectTF [14] and the Universal Protein Resource (UniProt) [15]. CollectTF provides well described and characterized, *in vivo* validated, TFs while UniProt is a comprehensive resource for protein sequence and annotation data. We used BacTFDB to train a deep learning model to predict new TFs and their families in genomes and metagenomes. Five model organisms *(Escherichia coli, Bacillus subtillis, Pseudomonas fluorescens, Azotobacter vinelandii and Caulobacter crescentus)* were used to test the performance and accuracy of PredicTF. We used the same approach to predict TFs from a clinical isolate *(P. aeruginosa* PAO1) and a metagenome sample isolated from an anaerobic ammonium oxidation community. We also determined if the predicted TFs were expressed in transcriptomes (isolate) and metatranscriptomes (microbial community), respectively (Fig. 1).

**Fig. 1.**
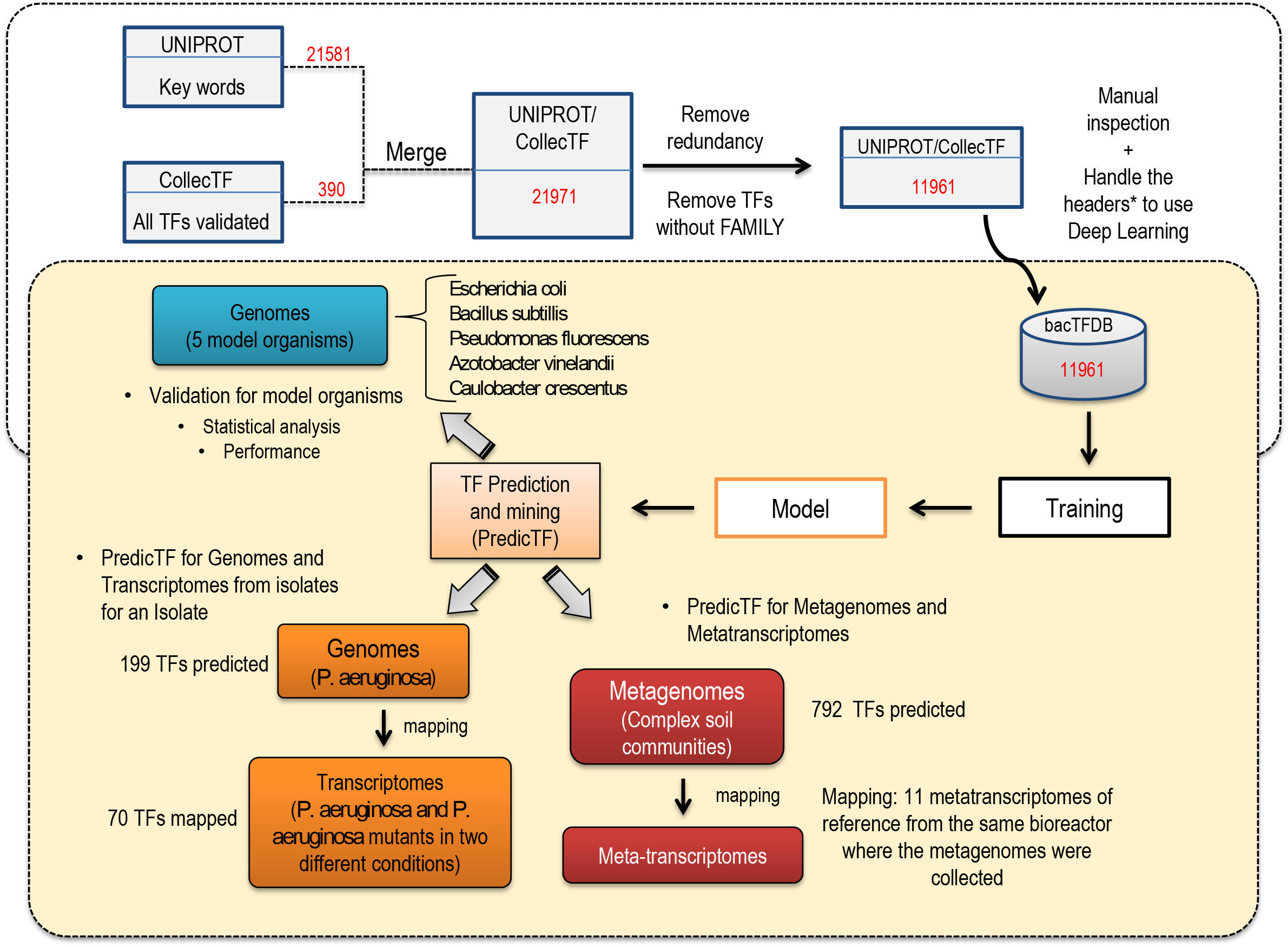
PredicTF workflow and testing. We collected publicly available data on TFs from two different databases: CollecTF and UNIPROT. After removing redundancies and filtering TFs well characterized, this data (BacTFDB) was used to train a deep learning model to predict new TFs and their families. Five model organisms (*Escherichia coli, Bacillus subtillis, Pseudomonas fluorescens, Azotobacter vinelandii and Caulobacter crescents*) were used to test the accuracy of PredicTF. Later, we used the same approach to predict TFs from an isolate (*P. aeruginosa*) and mapped TFs predicted in transcriptomics data (*P. aeruginosa* and mutants in two experimental conditions). Finally, we used our tool to predict TF for complex communities (metagenome) and mapped these TFs in their respective meta-transcriptomes.

### Database

BacTFDB is a robust and versatile bacterial TF database, it contains 11.691 TFs amino acid sequences spanning 1049 TF families and 720 different bacterial species. Fig. 2 shows the database distribution based on TF families and regulatory elements (Fig. 2A) and the distribution based on bacterial species (Fig. 2B). Although BacTFDB is composed by 11.961 TFs elements from 1049 different families and 720 organism’s species, Fig. 2 shows TFs families and species that accumulate more than 50 sequences. We will update BacTFDB annually by adding novel entries deposited in UniProt and CollecTF. BacTFDB was used in PredicTF’s deep learning model training. This model was later used to predict new TFs and their families in genomes and metagenomes.

**Fig. 2.**
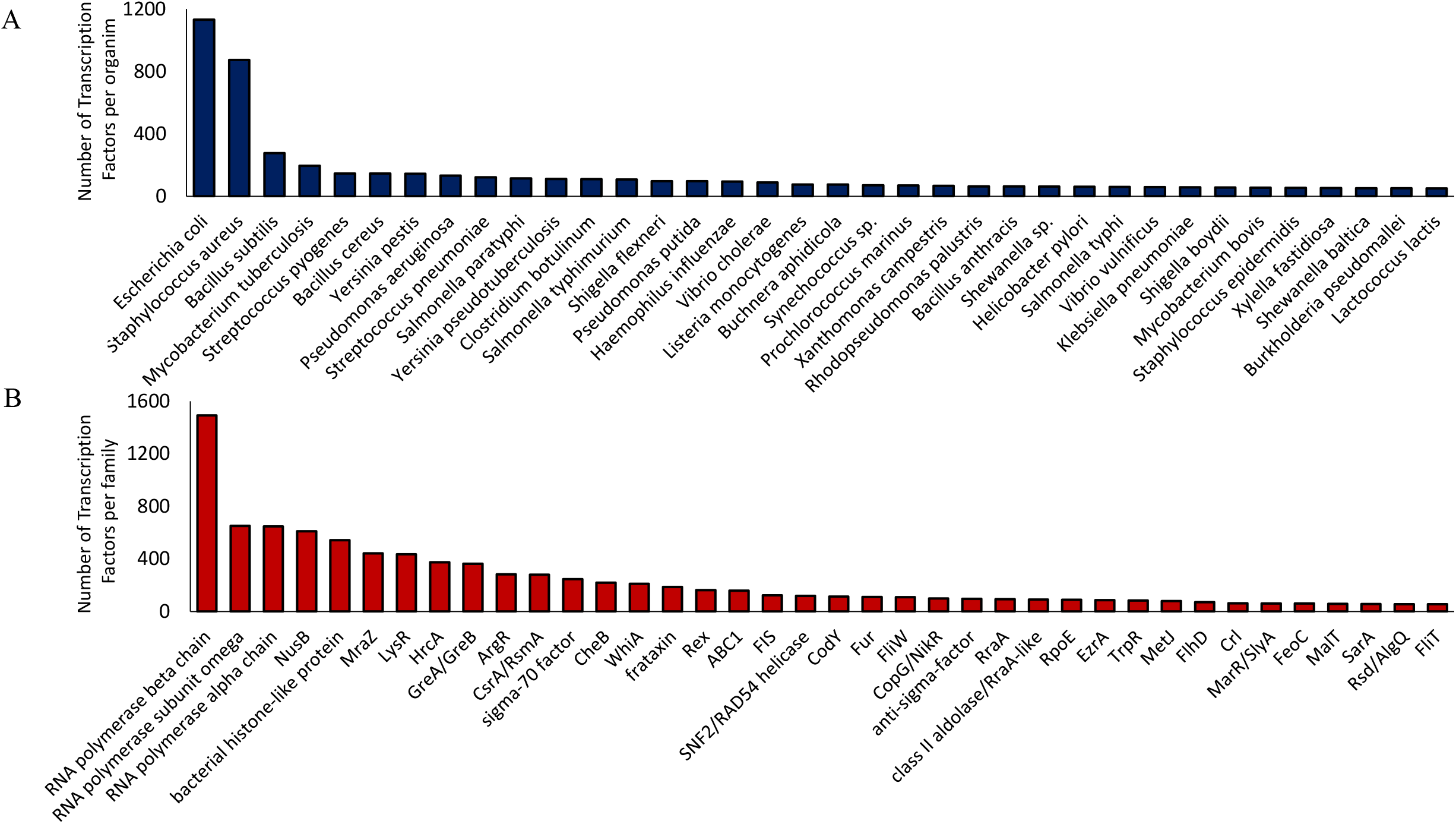
Database composition: Transcription Factor Database (BacTFDB) distribution. **A)** Database distribution based on the TFs and **B)** Regulatory Elements families and Organisms species. In these graphics only families with up to 50 sequences and only organisms that contributed with more than 50 sequences are shown.

### Performance and Accuracy

The performance and accuracy of PredicTF were evaluated through the prediction of TFs in five model organisms *(E. coli, B. subtillis, P. fluorescens, A. vinelandii and C. crescentus).* For each model organism a different PredicTF model was trained to predict TFs from full protein-length sequences (described in the implementation section).

The performance of PredicTF to identify TFs in the different model organisms ranged from 27% to 60% of the proteins described as TFs in the genomes of model organisms and the accuracy for experimentally validated TFs ranged from 73.91% and 91.43% (Table 1). Further, PredicTF was able to identify putative annotated TFs in the genomes of *E. coli* and *B. subtillis* with accuracies 85.71% and 100%, respectively (Table 1). No novel TF was predicted in the genome of *C. crescentus, P. fluorescens* and *A.vinelandii* (Table 1). TFs predicted by PredicTF for each organism, sorted by TF family, are shown in Fig. 3. For all organisms tested the most predicted TF family was LysR followed by OmpR/PhoB. The degree of accuracy obtained by PredicTF suggests that the deep learning strategy used is promising for the prediction of TFs in genomic or metagenomic data of bacterial species. PredicTF performance and accuracy can be further improved by expanding the number and diversity of sequences present in BacTFDB. As BacTFDB will be update yearly, we expect an improvement in TF identification of with every update.

**Fig. 3.**
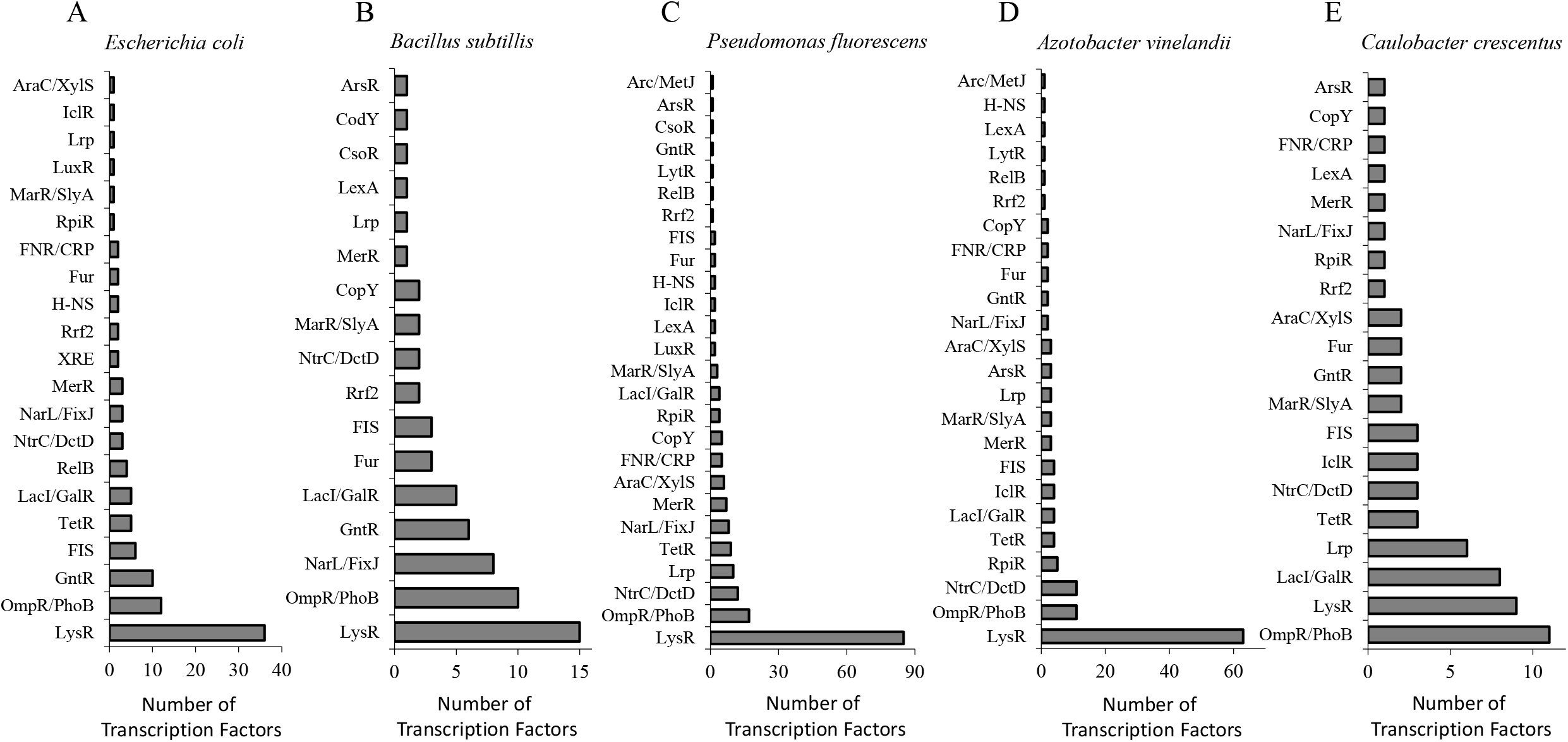
Prediction of TFs by PredicTF for genomes of model organisms. Prediction of TFs or 5 model organisms sorted by family. **A)** *Escherichia coli* **B)** *Bacillus subtillis* **C)** *Caulobacter crescentus* **D)** *Pseudomonas fluorescens* **E)** *Azotobacter vinelandii*

**Table 1.** PredicTF performance, accuracy for experimentally validated Transcription Factors (Accuracy EV) and accuracy for putative Transcription Factors (Accuracy PU) in genomes of model organisms.

| Organism | Performance<sup>a</sup> | Accuracy EV<sup>b</sup> | Accuracy PU<sup>c</sup> |
| --- | --- | --- | --- |
| *E. coli k12* | 35.40% | 88.51% | 85.71% |
| *B. subtilis* | 27.23% | 73.91% | 100% |
| *C. crescentus* | 38.04% | 83.93% | -<sup>d</sup> |
| *P. fluorescens* | 51.19% | 91.43% | - |
| *A. vinelandii* | 60.53% | 90.40% | - |

<sup>a</sup>Performance was calculated by the ratio of the total number of TFs predicted by PredicTF (*Predicted TFs*) to the total number of proteins annotated as TFs in NCBI (*Annotated TFs*) multiplied by 100;
<sup>b</sup>Accuracy EV was determined by the ratio of the total number of TFs predicted by PredicTF in agreement with NCBI annotation (*TFs predicted correctly*) to the total number of TFs predicted by PredicTF (*TFs predicted*) multiplied by 100;
<sup>c</sup>Accuracy TU was determined by the total number of putative TFs predicted correctly divided by putative TFs predicted multiplied by 100; *Putative TFs predicted correctly* is the total number of putative TFs predicted correctly by PredicTF in agreement with NCBI annotation; and, *Putative TFs predicted* is the total number of putative TFs predicted by PredicTF;
<sup>d</sup>Currently there are no putative annotated TFs described in the genome of *C. crescentus*, *P. fluorescens* and *A. vinelandii*

### Mining and Predicting TFs in Genomes and Transcriptomes from a bacterial isolate using PredicTF

PredicTF was used to predict TFs on the genome of *P. aeruginosa* PAO1 and these TFs were mapped in transcriptomes from the same isolate [16]. PredicTF predicted a total of 199 TFs in the *P. aeruginosa* PAO1 genome shown in Additional file 1: Fig. S1A by a family’s distribution graphic. These 199 TFs were mapped in the transcriptomic data of a reference of *P. aeruginosa* PAO1. Initially, the mapping was done in the transcriptome of *P. aeruginosa* PAO1 cultured in LB media. Using this strategy, we were able to map 69 of the 199 predicted TFs to the transcriptomes under the experimental conditions carried out by Hwang & Yoon, 2019 (Additional file 1: Fig. S1B) [16]. Next, the mappings were done for another three clinical mutants of *P. aeruginosa* PAO1 (Y82, Y71, Y89) cultured in LB media (absence of an antibiotic cocktail) (Additional file 2: Fig. S2A, S2C and S2F). The TFs family’s distribution for each *P. aeruginosa* PAO1 mutant cultured in presence of antibiotic cocktail is shown in the supplementary data (Additional file 2: Fig. S2B, S2D and S2F). These results demonstrate the potential of PredicTF in mapping regulatory elements in bacterial genomes and the use of this tool to map and compare TFs profiles after under different environmental conditions.

### Mining and Predicting TFs in a Metagenome and Metatranscriptome using PredicTF

PredicTF was used to profile TFs in one metagenome recovered from an anaerobic ammonium oxidation community [17] followed by the mapping of the predicted TFs in metatranscriptomes recovered from the same community (metatranscriptomes accession numbers can be found in Additional file 3: Table S1). A total of 792 TFs (Fig. 4A) were predicted in LAC_MetaG_1, an anaerobic ammonium oxidizing microbial community from an anammox membrane bioreactor [17]. These 792 TFs are distributed across 27 TF families (Fig. 4A) and are related to the regulation of functions such as the oxygen limitation response and late symbiotic functions (NarL/FixJ), phosphate regulon response (OmpR/PhoB), transcriptional activator for nitrogen-regulated promoters (NtrC/DctD) and ferric uptake regulation (Fur). To determine how a traditional annotation pipeline identify potential TF we used Prokka [18]. This tool was able to identify 1815 ORFs (Additional file 4: Table S2). PredicTF can be used with no previous knowledge regarding transcription factors, it is fast and it requires low memory when compared to Blast based annotation and it indicates only results of TFs with a specific TF family annotation. On the other hand, to identify TFs using Prokka one would need specialized training to mine the general annotation. Therefore, scientists with general microbiology background may take a long time to undergo this task. Further, Prokka gives no indication to the TF families of the putative annotated TFs. Time is also a drawback of using Prokka to mine TFs, we calculated we needed over 400 h to perform mine one single metagenomics library; in comparison, PredicTF needed 2 h to identify TF in the same metagenomics library.

**Fig. 4.**
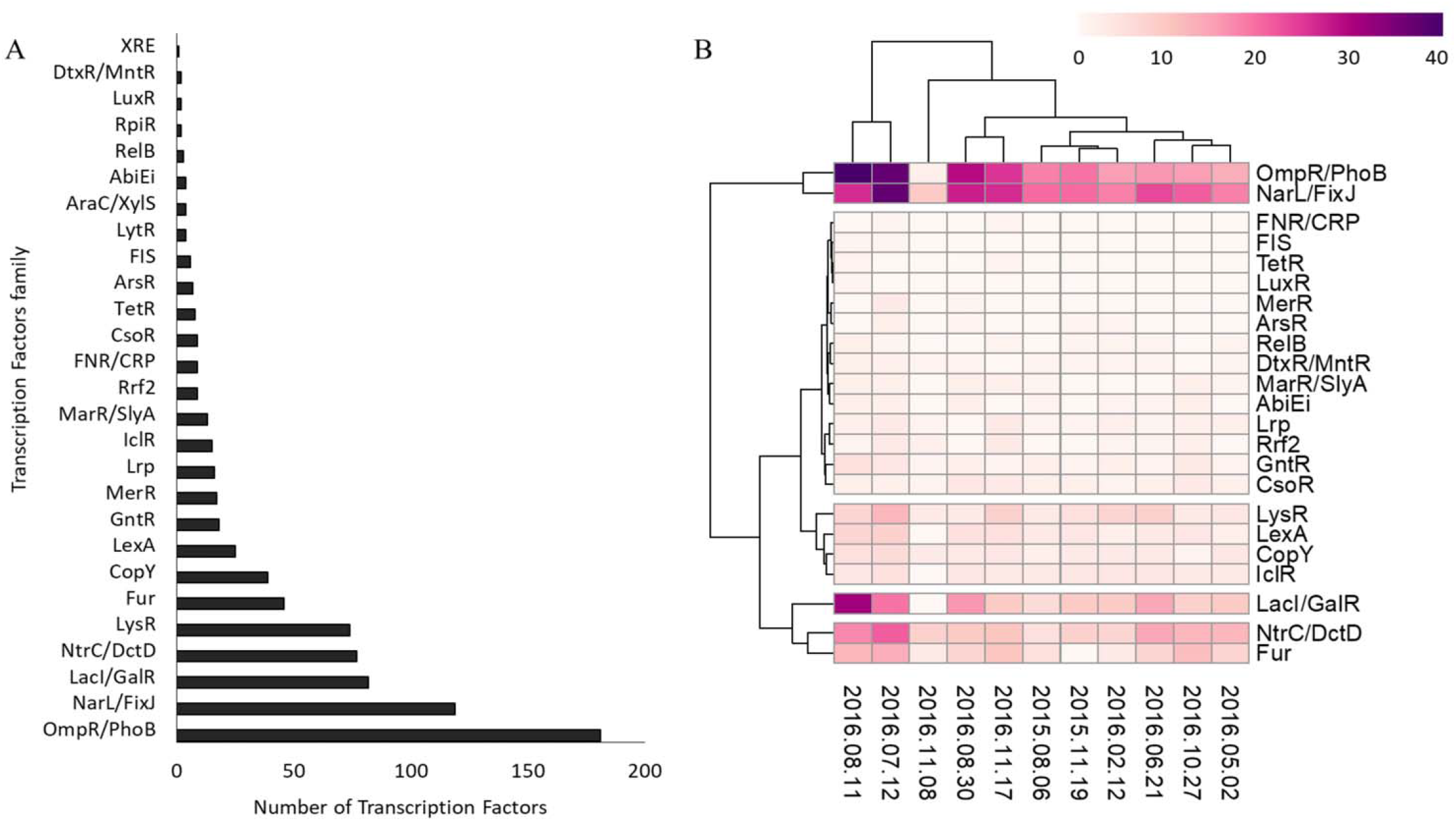
Recovery of novel Transcription Factors in one metagenome and eleven metatranscriptomes. **A)** PredicTF predicted 792 TFs were predicted in one anaerobic ammonium oxidizing microbial communities from anammox membrane bioreactor (LAC_MetaG_1) and were grouped by family. **B)** Using 792 TFs predicted in one metagenome, we mapped these TFs for 11 metatranscriptomes of reference from the same bioreactor where the metagenome was recovered.

Next, the 792 TFs were mapped in 11 metatranscriptomes collected in different dates from the same bioreactor where the metagenome was recovered (Additional file 5: Table S3, Fig. 4B). Clustering analysis demonstrated the presence of five different groups of TFs families based on the number of transcription factor families expressed in each library (Fig. 4B). It is interesting to note that the two most abundant clusters in the heatmap are directly related to the oxygen limitation caused by the anaerobic ammonium oxidizing cultivation. In a bioreactor where oxygen is limited, an increase in the amount of nitrogen and phosphate is expected. The presence of N and P diverts the metabolism of the microbial community towards the production of regulators (TFs) that help to maintain community stability. Clustering analyzes can be helpful to demonstrate the similarity between metatranscriptomic libraries based on the occurrence of TFs. This strategy can be useful to compare the profiles of TFs expressed in different environmental situations (comparing libraries with different metadata) creating patterns of TFs expression. Exploration of TF profiling in microbial communities (metagenomes or metatranscriptomes) will allow the exploration of regulation within complex microbial communities. Further, The recovery of metagenome assembled genomes is becoming standard in metagenomics studies [19–21]. The use of PredicTF together with the recovery of metagenome assembled genomes will allow the exploration of species-specific molecular mechanisms involved in the regulation of different ecosystem processes.

### Conclusions

A better understanding of TFs in a bacterial community context open revenue for the exploration of gene regulation in ecosystems where bacteria play a key role. Our deep learning strategy was based on a novel and robust TF bacterial database (BacTFDB) with over 11 thousand TFs and their respective families. BacTFDB is a unique resource for the exploration of TFs and it provided the data to train a model within PredicTF capable of predicting novel TFs from genomes and metagenomes. PredicTF is the first pipeline designed to predict and annotate TFs in complex microbial communities. The prediction of TFs can provide information for those aiming to study and understand bacterial communities within a context of gene regulation. We also demonstrated that PredicTF can be used to predict novel TFs in metagenomes and metatrascriptomes creating the potential profile for regulatory elements in complex microbial communities.

PredicTF is a flexible open source pipeline able to predict and annotate TFs in genomes and metagenomes and can be found at https://github.com/mdsufz/PredicTF.

## Methods

### BacTFDB - Bacterial Transcription Factor Data Base

To create a novel Bacterial Transcription Factor Data Base (BacTFDB), we collected data from two publicly available databases. Initially, we chose to collect data from CollecTF [14], a well described and characterized database. Since CollecTF does not provide an application programming interface (API) for bulk download, we developed a Python code (version 2.7) using the Beautiful Soup 4.4.0 library to recover the data from CollecTF. With this strategy we listed 390 TF experimentally validated amino acid sequences distributed over 44 TF families. The script can be found at https://github.com/mdsufz/PredicTF.

Additionally, we retrieved TF amino acid sequences from UniProt using UniProt’s API. We downloaded sequences of interest by adding a filter with the key words (Transcription factor, transcriptional factor, regulator, transcriptional repressor, transcriptional activator, transcriptional regulator). After, we filtered for Reviewed (Swiss-Prot) - Manually annotated sequences that belonged to the bacteria taxonomy. The UniProt API was accessed on 8^th^ September-2019 and a total of 21.581 TF amino acid sequences, with applied filters, were collected. We merged the data collected from CollecTF and UniProt databases which resulted in a total of 21.971 TFs. Next, we removed redundant TF entries and TF sequences lacking a TF family since PredicTF was designed to also assign TF family. Finally, a manual inspection was performed to remove case sensitive and presence of characters associated to the database header. The first version of BacTFDB contains a total of 11.691 unique TF sequences. A summary of the information contained in BacTFDB can be found in the supplementary data (Additional file 6: Fig. S4). To evaluate PredicTF in model organisms we created 5 subsets of BacTFDB. The description of these subsets can be found in the supplementary data (Additional file 7: Table S4).

### Mapping Transcription Factors using PredictTF

We used a deep learning approach similar to that found in DeepARG [22]. Supervised machine learning models are usually divided into characterization, training, and prediction units. Briefly, our approach uses the concept of dissimilarity-based classification [23] where sequences are represented and featured by their sequence similarity to known genes. BacTFDB was used to train and test the deep learning model (https://github.com/mdsufz/PredicTF) and latter validated in model organisms. Next, PredicTF was used to predict novel TFs from full protein-length sequences in genomes and in one metagenome. After prediction, the data was mapped in transcriptomes and metatranscriptomes from samples where the genetic potential was determined.

Using PredicTF, we trained five different models – one for each model organism (Additional file 3: Table S1). For each model, the TFs affiliated with the respective model organism were removed prior to training to avoid overfitting. PredicTF-no-coli was trained to predict TFs in *E. coli,* PredicTF-no-subtilis was trained to predict TFs in *B. subtilis,* PredicTF-no-crescentus was trained to predict TFs in *C. crescentus*, PredicTF-no-fluorescens was trained to predict TFs in *P. fluorescens* and PredicTF-no-vinelandii was trained to predict TFs in *A. vinelandii*.

### Performance and accuracy calculation

We evaluated PredicTF by calculating accuracy and performance. Performance can be deemed to be the fulfillment of a task. In PredicTF case, performance is how good TF predictions are. Using model organisms (see later in the session *Prediction of Transcription Factors in model organisms*), performance was calculated by quantifying the number of TFs that PredicTF was able to predict divided by number of TFs already described and annotated for our model organisms (Additional file 7: Equation 1). Accuracy indicates how correct the predictions performed by PredicTF are. Also using data of model organism, accuracy was determined by calculating the number of TFs correctly predicted divided by the total number of TFs predicted by PredicTF. We divided accuracy in two categories. In the first accuracy category, we determined accuracy against experimentally validated TFs (Additional file 7: Equation 2). In the second accuracy category, we determined accuracy against TFs without experimental validation (Additional file 7: Equation 3); i.e., putative TFs. The performance, accuracy, and accuracy for putative TFs were calculated as the ratio of predicted to annotated TFs. Accuracy was quantified as the fraction of correctly predicted TFs among all predictions.

### Prediction of Transcription Factors in model organisms

We selected bacterial species that have been widely studied as model organisms. Some bacterial species became model organisms for TF studies because they are easy to maintain and grow in a laboratory setting and to manipulate in pure culture experiments. Five complete genomes from model organisms (*E. coli, B. subtillis, P. fluorescens, A. vinelandii and C. crescentus*) were downloaded directly from NCBI. The strains details and accession number (RefSeq) for all selected organisms are listed in the supplementary data (Additional file 3: Table S1). By evaluating PredicTF using model organisms (Additional file 6: Table S3) we extrapolated performance and accuracy of our deep learn model. Since known TFs for each organism were removed from each the training dataset, we eliminate the possibility of mapping TFs already known and annotated for each of the different species. Performance, accuracy and accuracy for putative TFs of PredicTF for these five model organisms were calculated using Equations 1, 2 and 3.

### Prediction of Transcription Factors in a clinical isolate

We demonstrated the use of PredicTF in a previously sequenced *P. aeruginosa* (PAO1) genome, a clinical isolate publicly available in NCBI (accession number **NC_002516.2**). *P. aeruginosa* PAO1 was selected because its genome has been sequenced and because of the availability of transcriptomes from three clinical mutants of PAO1 (Y71, Y82, and Y89) grown in the presence and absence of an antibiotic cocktail. The transcriptomes of *P. aeruginosa* PAO1 mutants Y71, Y82, and Y89 are available in NCBI (Bioproject identifier **PRJNA479711**) [16]. These clinical *P. aeruginosa* PAO1 mutants were isolated from the sputa of three different pneumonia patients. Transcriptomes of *P. aeruginosa* PAO1 wild type and its mutants cultured in two different conditions (LB medium and LB medium in presence of antibiotic cocktail) have been previously described [16]. We used this data to determine the TF profile in these *P. aeruginosa* PAO1 mutants grown in two different conditions.

PredicTF was first used to predict TFs in the *P. aeruginosa* PAO1 genome. Next, the predicted TFs were mapped to the transcriptomes of the *P. aeruginosa* PAO1 mutants Y71, Y82 and Y89 (see later). Further description of the mapping of the transcriptomes to the genomes is available at https://github.com/mdsufz/PredicTF. The PredicTF model used in this step was trained with the full database BacTFDB. All accession numbers used in this work are listed in the supplementary data (Additional file 3: Table S1).

### Prediction of Transcription Factors in Complex Microbial Communities

To test PredicTF in a complex microbial community, we used an anaerobic ammonium oxidizing (anammox) microbial community from an anammox membrane bioreactor metagenome (LAC_MetaG_1) (data publicly available at NCBI bioproject via accession number **PRJNA511011**) [17]. We removed short and low-quality reads using Trim Galore - v0.0.4 dev according developer’s instructions [24]. Over 50 million reads survived this step and were assembled using the *de novo* assembler metaSPADES - v3.12.0 [25]. The assembly was translated from nucleotide to amino acid sequences, considering all possible translation frames, using emboss transeq [26]. The translated assembly was then used as input for the prediction of transcription factors using PredicTF. The region from each predicted TF was extracted. These putative TFs were later used in the mapping TFs to metatranscriptomes.

We checked if the putative TFs predicted in the metagenomes were transcribed by checking if the metatranscriptomic libraries were mapping to those regions. The metatranscriptomic and metagenomic libraries used in this step belonged to the same bioreactor. These metatranscriptomes are publicly available at the European Nucleotide Archive under the accession numbers SRR7091385, SRR7523233, SRR7523244, SRR7523245, SRR7091400, SRR7091401, SRR7091381, SRR7091402, SRR7091406, SRR7523243, SRR7523246. These 11 metatranscriptomes were used to demonstrate the effectiveness of the pipeline and to indicate the potential of PredicTF to profile transcription factors in complex microbial communities. All accession numbers used in this work are listed in the supplementary data (Additional file 3: Table S1).

To have a baseline comparison with a traditional annotation pipeline, we used Prokka [18] to annotate the same anammox membrane bioreactor metagenome (LAC_MetaG_1). We mined the annotation by hand with specialized knowledge of scientists specialized in Transcription Factors. We did not determine the families as this work would need to be done for every single hit individually using the output of Prokka.

### Mapping transcription factors to transcriptomes and metatranscriptomes

Each transcriptomic and metatranscriptomic library was quality controlled by removing short and low-quality reads using Trim Galore - v0.0.4 dev [24]. The 7 transcriptomic libraries for the *P. aeruginosa* PAO1 wild type and mutants showed at least 26 million paired end reads after quality checking. The 11 metatranscriptomic libraries yielded over 50 million reads per library after quality check. After, the remaining transcriptomic and metatranscriptomic reads were mapped to their respective assembled genome or metagenome using Bowtie2 - v2.3.0 [27]. The number of reads mapped, and the regions covered was extracted using SAMTools - v1.9 [28] and python 2.7. The regions of the genome or metagenome assembly covered by transcriptomic or metatranscriptomic reads were then crossed-referenced with the regions of their respective assembly which PredicTF assigned as putative TFs creating a TF profile for each transcript and metatranscriptome. A detailed description on the mapping of RNA-seq data to their respective genome or metagenome assembly can be found at the PredicTF github (https://github.com/mdsufz/PredicTF).

## List of Abbreviations

(TFs): Transcription Factors
(BacTFDB): Bacterial Transcription Factor Data Base
(TFBSs): Transcription factor binding sites
(anammox): anaerobic ammonium oxidizing

## Declaration Sections

### Ethics approval and consent to participate

Not applicable

### Consent for publication

Not applicable

### Availability of data and materials

Project name: PredicTF

Project home page: https://github.com/mdsufz/PredicTF

Operating system: Linux64

Programming languages: Python 2.7

Other requirements: DIAMOND [29]; Nolearn Lasagne deep learning library [30]; Sklearn machine learning routines (https://scikit-learn.org/stable/) [31]; Theano (http://deeplearning.net/software/theano/) [32]. Trim Galore - v0.0.4 dev (https://www.bioinformatics.babraham.ac.uk/projects/trim_galore/) [24]. MetaSPADES - v3.12.0 (https://github.com/ablab/spades#meta) [25]. Emboss transeq (http://www.bioinformatics.nl/cgi-bin/emboss/transeq) [26]. Bowtie2 - v2.3.0 (https://sourceforge.net/projects/bowtie-bio/) [27]. SAMTools - v1.9 (http://github.com/samtools/) [28].

Genomes of the model organisms used in the Tool Validation step are available at the National Center for Biotechnology Information (https://www.ncbi.nlm.nih.gov/) under the accession numbers NC_000913.3, NC_000964.3, NC_011916.1, NC_021149.1, and NC_016830. The datasets supporting the Prediction of Transcription Factors in a clinical isolate of this article are available at National Center for Biotechnology Information (https://www.ncbi.nlm.nih.gov/) under the accession number NC_002516.2 (genome) and study accession PRJNA479711 (transcriptomes). The datasets used for the Prediction of Transcription Factors in Complex Microbial Communities of this study are available at National Center for Biotechnology Information (https://www.ncbi.nlm.nih.gov/) under the study accession PRJNA511011. The respective data sets of metatranscriptomes used are available at National Center for Biotechnology Information (https://www.ncbi.nlm.nih.gov/) under the SRA numbers SRR7091385, SRR7523233, SRR7523244, SRR7523245, SRR7091400, SRR7091401, SRR7091381, SRR7091402, SRR7091406, SRR7523243, SRR7523246 and the Joint Genome Institute (https://jgi.doe.gov/) under the Gold Analysis Project identifiers Gp0267156, Gp0267150, Gp0267154, Gp0267155, Gp0267157, Gp0267158, Gp026715, Gp0267159, Gp0267152, Gp0267153, Gp0267160. All analysis, results and scripts used to generate figures are available at https://github.com/mdsufz/PredicTF.

### Authors’ contributions

LMOM, PFS, RSR, and UNR developed the concept of PredicTF. LMOM, JS, UNR developed the PredicTF workflow. LMOM, JS, and UNR performed the benchmarks. LMOM provided information and data for the creation BacTFDB dataset. RBT and UNR performed the metagenome and metatranscriptome analysis. LMOM and UNR wrote the manuscript. All authors read and approved the manuscript.

## Additional files

**Additional file 1: Fig. S1**

Transcription factor (TF) families predicted for *Pseudomonas aeruginosa* PAO1 genome (accession number NC_002516.2) [18] using PredicTF and their mapping to *P. aeruginosa* PAO1 growing in LB medium. A) A total of 199 TFs distributed in 25 TF families were predicted in the *P. aeruginosa* PAO1 genome. B) These 199 TFs were mapped in the transcriptomic data of a reference of *P. aeruginosa* PAO1 (Bioproject identifier PRJNA479711) [18]. Initially, the mapping was done in the transcriptome of *P. aeruginosa* PAO1 cultured in LB media. Using this strategy, we were able to map 69 of the 199 predicted TFs to the transcriptome.

**Additional file 2: Fig. S2**

Transcription Factor (TF) family profiles in three *Pseudomonas aeruginosa* PAO1 mutants. After the prediction of Transcription Factors (TFs) using *P. aeruginosa* PAO1 genome, we mapped transcriptomes from three *P. aeruginosa* PAO1 mutants (Y82, Y71, Y89) cultured in LB media (A, C and F). After, the mapping was done for each *P. aeruginosa* PAO1 mutant cultured in presence of antibiotic cocktail (B, D and E). *P. aeruginosa* PAO1 mutant Y82 (A, B); P. aeruginosa PAO1 mutant Y71 (C, D); *P. aeruginosa* PAO1 mutant Y89 (E, F).

**Additional file 3: Table S1**

Accession number for 5 model organisms, *Pseudomonas aeruginosa* PAO1 genome and transcriptomes and Complex Microbial Communities used to validate and test PredicTF.

**Additional file 4: Table S2**

Transcription factors from the metagenome of an anaerobic ammonium oxidizing microbial community from an anammox membrane bioreactor (LAC_MetaG_1) mined and hand curated from a general annotation generated using Prokka.

**Additional file 5: Table S3**

Number of Transcription Factors (TFs) per TF family mapped to each of the 11 metatranscriptomes of reference from the same bioreactor where the metagenome (accession number PRJNA511011, NCBI) used to predict the putative TFs was collected. The different metatranscriptomes are represented by their European Nucleotide Archive accession numbers.

**Additional file 6: Fig. S4**

Bacterial Transcription Factor Data Base (BacTFDB) were created from from two publicly available databases. We collect 390 TFs from CollecTF and 21.581 from UniProt (accessed 8-Sep-2019) accumulating 21.581 Transcription Factor (TF) amino acid sequences. We merged the data from CollecTF and UniProt databases resulting in a total of 21.971 TFs amino acid. We removed redundant TF entries and since PredicTF was designed to also assign TF family, TF sequences lacking a TF family were removed. Finally, a manual inspection was performed to remove misleading of spelling, case sensitive and presence of characters associate to the database header. The final database (BacTFDB) contains a total of 11.691 TF unique sequences.

**Additional file 7: Table S4**

Description of the bacterial transcriptional factors database (BacTFDB) subsets used to train models to predict Trancription Factors in model organisms

**Additional file 8**

Equations used to calculate PredicTF’s accuracy and performance.

