## Additional_File_1 for "PredicTF: a tool to predict bacterial transcription factors in complex microbial communities"

---

### Additional file 1: Fig. S1

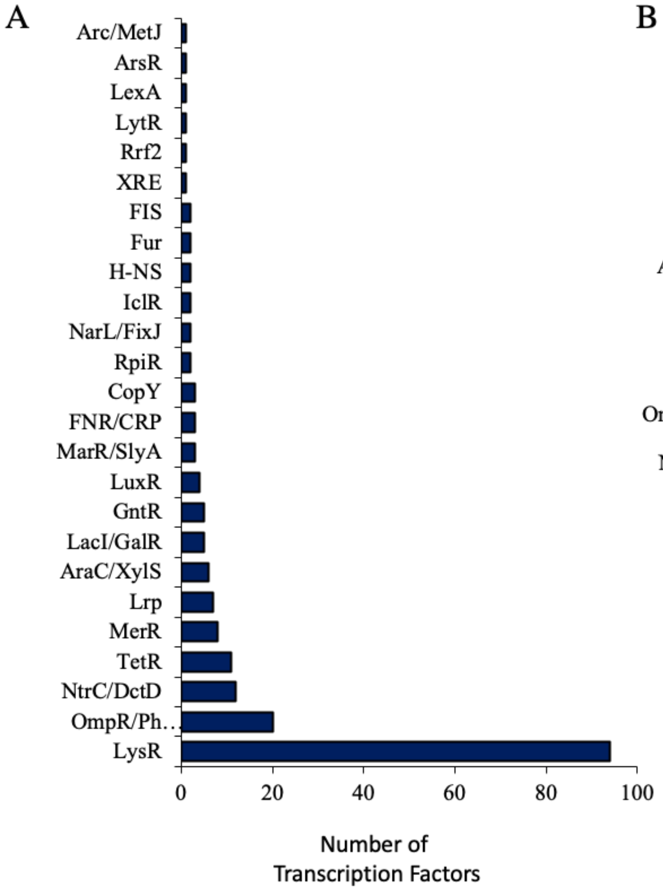

**PredicTF was used for the prediction of Transcription Factors (TFs) using *Pseudomonas aeruginosa* PAO1 genome** A) the graphic sorted by the family's distribution shows a total of 199 TFs that were predicted in the *P. aeruginosa* PAO1 genome. B) These 199 TFs were mapped in the transcriptomic data of a reference of *P. aeruginosa* PAO1. Initially, the mapping was done in the transcriptome of *P. aeruginosa* PAO1 cultured in LB media. Using this strategy, we were able to map 69 of the 199 predicted TFs to the transcriptome.
