## Additional_File_2 for "PredicTF: a tool to predict bacterial transcription factors in complex microbial communities"

### Additional file 2: Fig. S2

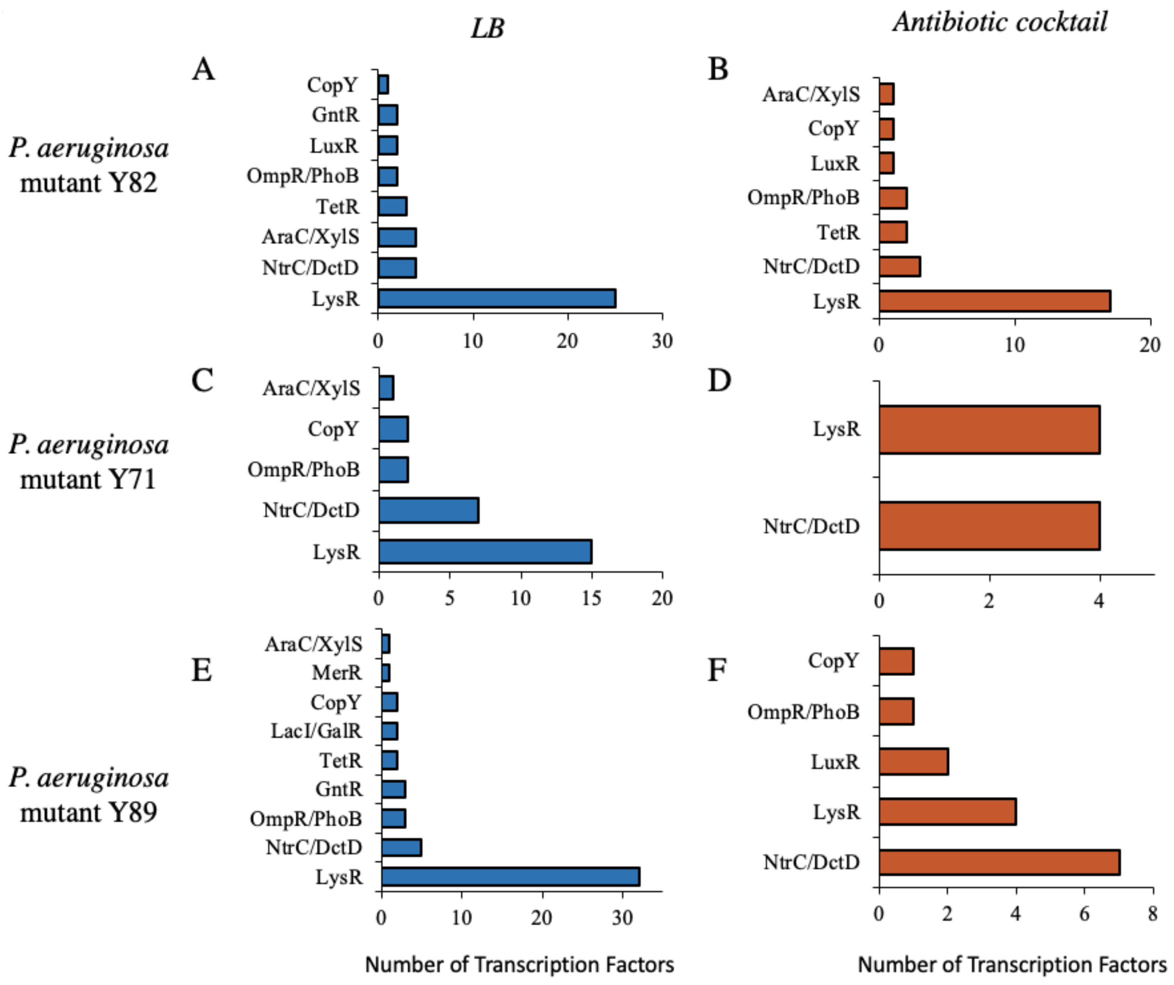


**Legend.** Transcription Factor (TF) family profiles in three *Pseudomonas aeruginosa* PAO1 mutants. After the prediction of Transcription Factors (TFs) using *Pseudomonas aeruginosa* PAO1 genome, we mapped transcriptomes from three *P. aeruginosa* PAO1 mutants (Y82, Y71, Y89) cultured in LB media (A, C and F). After, the mapping was done for each *P. aeruginosa* PAO1 mutant cultured in presence of antibiotic cocktail (B, D and E). *P. aeruginosa* PAO1 mutant Y82 (A, B); *P. aeruginosa* PAO1 mutant Y71 (C, D); *P. aeruginosa* PAO1 mutant Y89 (E, F).
