## Additional_File_3 for "PredicTF: a tool to predict bacterial transcription factors in complex microbial communities"

---

**Additional file 3: Table S1** Accession number for 5 model organisms, *Pseudomonas aeruginosa* PAO1 genome and transcriptomes and Complex Microbial Communities validated with PredicTF.

| *Model Organisms* | | |
|---|---|---|
| **Description** | **RefSeq** | **Location** |
| ***Escherichia coli* (strain K-12 sub strain MG1655)** | NC_000913.3 | (NCBI) Genbank |
| ***Bacillus subtilis* (strain 168)** | NC_000964.3 | (NCBI) Genbank |
| ***Caulobacter vibrioides* (strain NA1000 / CB15N) (*Caulobacter crescentus*)** | NC_011916.1 | (NCBI) Genbank |
| ***Azotobacter vinelandii* (strain DJ / ATCC BAA-1303)** | NC_021149.1 | (NCBI) Genbank |
| ***Pseudomonas fluorescens* (strain F113)** | NC_016830.1 | (NCBI) Genbank |
| ***Clinical Isolate*** | | |
| **Description** | **RefSeq** | **Location** |

| | | |
|---|---|---|
| ***Pseudomonas aeruginosa* (PAO1) genome** | NC_002516.2 | (NCBI) Genbank |
| **Transcriptomes from three clinical isolates (Y71, Y82, and Y89)** | PRJNA479711 | (NCBI) Genbank |

***Complex Microbial Communities***

| **Description** | **RefSeq** | **Identification** | **Date of Collection** | **Location** |
|---|---|---|---|---|
| **Metagenomic dataset** | PRJNA511011 | LAC_MetaG_1 | | (NCBI) Genbank |
| **Meta-transcriptomes of anaerobic ammonium oxidizing microbial communities from anammox membrane bioreactor (MBR)** | SRR7091385 | LAC_MetaT_1 | 2015-08-06 | European Nucleotide Archive (ENA) |
| | SRR7523233 | LAC_MetaT_2 | 2015-11-19 | |
| | SRR7523244 | LAC_MetaT_3 | 2016-02-12 | |
| | SRR7523245 | LAC_MetaT_4 | 2016-05-02 | |
| | SRR7091400 | LAC_MetaT_5 | 2016-06-21 | |
| | SRR7091401 | LAC_MetaT_6 | 2016-07-12 | |
| | SRR7091381 | LAC_MetaT_7 | 2016-08-11 | |
| | SRR7091402 | LAC_MetaT_8 | 2016-08-30 | |
| | SRR7091406 | LAC_MetaT_9 | 2016-10-27 | |
| | SRR7523243 | LAC_MetaT_10 | 2016-11-08 | |
| | SRR7523246 | LAC_MetaT_11 | 2016-11-17 | |
