## Additional_File_4 for "PredicTF: a tool to predict bacterial transcription factors in complex microbial communities"

**Additional file 7:**

**Table S2: Transcription factors from the metagenome of an anaerobic ammonium oxidizing microbial community from an anammox membrane bioreactor (LAC_MetaG_1) mined and hand curated from a general annotation generated using Prokka.**

| locus_tag | product |
|---|---|
| **KCPMINPF_00018** | Regulatory protein RecX |
| **KCPMINPF_00282** | Phosphate regulon transcriptional regulatory protein PhoB |
| **KCPMINPF_00285** | Transcriptional regulatory protein CusR |
| **KCPMINPF_00319** | Transcriptional regulatory protein OmpR |
| **KCPMINPF_00332** | Regulatory protein LuxO |
| **KCPMINPF_00360** | DnaA regulatory inactivator Hda |
| **KCPMINPF_00389** | Hydrogen peroxide-inducible genes activator |
| **KCPMINPF_00667** | Copper-sensing transcriptional repressor CsoR |
| **KCPMINPF_00697** | Transcriptional regulatory protein QseB |
| **KCPMINPF_00700** | Transcriptional activator protein AnoR |
| **KCPMINPF_01226** | Nitrogen regulatory protein |
| **KCPMINPF_01534** | Transcriptional regulatory protein ZraR |
| **KCPMINPF_01666** | Bifunctional transcriptional activator/DNA repair enzyme Ada |
| **KCPMINPF_01678** | Heat-inducible transcription repressor HrcA |
| **KCPMINPF_01778** | Transcriptional repressor NrdR |

| | |
|---|---|
| **KCPMINPF_01864** | Transcriptional regulatory protein ZraR |
| **KCPMINPF_01871** | flagellum biosynthesis repressor protein FlbT |
| **KCPMINPF_01880** | RNA polymerase-binding transcription factor DksA |
| **KCPMINPF_02102** | Transcriptional activator protein CopR |
| **KCPMINPF_02113** | Redox-sensing transcriptional repressor Rex 1 |
| **KCPMINPF_02156** | KDP operon transcriptional regulatory protein KdpE |
| **KCPMINPF_02187** | Transcriptional regulatory protein LnrK |
| **KCPMINPF_02223** | DNA-binding transcriptional activator DevR/DosR |
| **KCPMINPF_02225** | Alkaline phosphatase synthesis transcriptional regulatory protein PhoP |
| **KCPMINPF_02284** | Regulatory protein AtoC |
| **KCPMINPF_02362** | Regulatory protein RecX |
| **KCPMINPF_02483** | Heat-inducible transcription repressor HrcA |
| **KCPMINPF_02687** | Transcriptional regulatory protein LiaR |
| **KCPMINPF_02701** | Transcriptional regulatory protein LnrK |
| **KCPMINPF_02805** | Ribose operon repressor |
| **KCPMINPF_02822** | Transcriptional regulatory protein WalR |
| **KCPMINPF_02855** | N-acetylglucosamine repressor |
| **KCPMINPF_02954** | Bifunctional transcriptional activator/DNA repair enzyme Ada |
| **KCPMINPF_02998** | DNA-binding transcriptional regulator BolA |
| **KCPMINPF_03057** | LexA repressor |
| **KCPMINPF_03096** | Sigma factor AlgU regulatory protein MucB |
| **KCPMINPF_03148** | Regulatory protein AtoC |
| **KCPMINPF_03226** | Transcriptional repressor NrdR |
| **KCPMINPF_03275** | Transcriptional regulatory protein BaeR |
| **KCPMINPF_03362** | Phosphate regulon transcriptional regulatory protein PhoB |
| **KCPMINPF_03435** | Regulatory protein AtoC |
| **KCPMINPF_03461** | DNA-binding transcriptional activator HyfR |
| **KCPMINPF_03514** | Pca regulon regulatory protein |
| **KCPMINPF_03553** | Phosphate regulon transcriptional regulatory protein PhoB |
| **KCPMINPF_03610** | Regulatory protein RecX |
| **KCPMINPF_03643** | Regulatory protein AtoC |
| **KCPMINPF_03670** | Transcriptional regulator MraZ |
| **KCPMINPF_03672** | Penicillin-binding protein activator LpoA |
| **KCPMINPF_03761** | Phosphate regulon transcriptional regulatory protein PhoB |
| **KCPMINPF_03788** | Transcriptional regulatory protein DegU |
| **KCPMINPF_03814** | Oxygen regulatory protein NreC |
| **KCPMINPF_03841** | Transcriptional regulatory protein LiaR |
| **KCPMINPF_03977** | Ribose operon repressor |
| **KCPMINPF_04134** | Alkaline phosphatase synthesis transcriptional regulatory protein PhoP |
| **KCPMINPF_04355** | Iron-dependent repressor IdeR |
| **KCPMINPF_04394** | Methanol dehydrogenase activator |
| **KCPMINPF_04549** | Transcriptional regulatory protein DegU |
| **KCPMINPF_04560** | Transcriptional regulatory protein LiaR |

| | |
|---|---|
| **KCPMINPF_04586** | RNA polymerase-binding transcription factor CarD |
| **KCPMINPF_04647** | LexA repressor |
| **KCPMINPF_04672** | DNA-binding transcriptional regulator NtrC |
| **KCPMINPF_04801** | Transcriptional regulatory protein DegU |
| **KCPMINPF_04820** | Copper-sensing transcriptional repressor CsoR |
| **KCPMINPF_04926** | Zinc-specific metallo-regulatory protein |
| **KCPMINPF_04974** | Transcriptional repressor NrdR |
| **KCPMINPF_05013** | Transcriptional regulatory protein AfsQ1 |
| **KCPMINPF_05015** | Alkaline phosphatase synthesis transcriptional regulatory protein SphR |
| **KCPMINPF_05050** | Transcriptional regulatory protein LiaR |
| **KCPMINPF_05125** | Heat-inducible transcription repressor HrcA |
| **KCPMINPF_05233** | Transcriptional regulatory protein DegU |
| **KCPMINPF_05321** | Regulatory protein AtoC |
| **KCPMINPF_05367** | Alkaline phosphatase synthesis transcriptional regulatory protein PhoP |
| **KCPMINPF_05405** | Leucine-responsive regulatory protein |
| **KCPMINPF_05586** | Alkaline phosphatase synthesis transcriptional regulatory protein PhoP |
| **KCPMINPF_05718** | Transcriptional regulator KdgR |
| **KCPMINPF_05762** | Regulatory protein RecX |
| **KCPMINPF_05970** | Transcriptional regulatory protein ComA |
| **KCPMINPF_06062** | Leucine-responsive regulatory protein |
| **KCPMINPF_06066** | DNA-binding transcriptional activator DecR |
| **KCPMINPF_06091** | Transcriptional activator protein NhaR |
| **KCPMINPF_06212** | Arginine repressor |
| **KCPMINPF_06233** | RNA polymerase-binding transcription factor DksA |
| **KCPMINPF_06252** | Nitrogen regulatory protein P-II |
| **KCPMINPF_06316** | Transcriptional regulatory protein ZraR |
| **KCPMINPF_06414** | DNA-binding transcriptional regulator NtrC |
| **KCPMINPF_06592** | Transcriptional regulator SlyA |
| **KCPMINPF_06679** | Heat-inducible transcription repressor HrcA |
| **KCPMINPF_06875** | Transcriptional regulatory protein DegU |
| **KCPMINPF_06907** | ATP phosphoribosyltransferase regulatory subunit |
| **KCPMINPF_06920** | Fumarate and nitrate reduction regulatory protein |
| **KCPMINPF_06922** | Transcriptional regulatory protein OmpR |
| **KCPMINPF_06982** | DNA-binding transcriptional regulator NtrC |
| **KCPMINPF_07122** | DNA-binding transcriptional regulator BolA |
| **KCPMINPF_07148** | Transcriptional regulator HilA |
| **KCPMINPF_07186** | Phosphoenolpyruvate synthase regulatory protein |
| **KCPMINPF_07242** | Transcriptional regulatory protein OmpR |
| **KCPMINPF_07483** | Transcriptional activator protein CopR |
| **KCPMINPF_07529** | Transcriptional regulator MtlR |
| **KCPMINPF_07659** | Transcriptional regulatory protein WalR |
| **KCPMINPF_07664** | Alkaline phosphatase synthesis transcriptional regulatory protein PhoP |
| **KCPMINPF_07670** | Transcriptional regulatory protein BaeR |

| | |
|---|---|
| **KCPMINPF_07839** | Transcriptional regulator LsrR |
| **KCPMINPF_07913** | Transcriptional regulatory protein QseB |
| **KCPMINPF_07929** | Photosynthetic apparatus regulatory protein RegA |
| **KCPMINPF_07947** | Transcriptional regulatory protein OmpR |
| **KCPMINPF_08029** | Nitrogen regulatory protein P-II |
| **KCPMINPF_08069** | Transcriptional activator HlyU |
| **KCPMINPF_08095** | Transcriptional activator protein Anr |
| **KCPMINPF_08265** | Transcriptional activatory protein AadR |
| **KCPMINPF_08293** | Transcriptional activator protein Anr |
| **KCPMINPF_08477** | Alkaline phosphatase synthesis transcriptional regulatory protein PhoP |
| **KCPMINPF_08914** | Regulatory protein RecX |
| **KCPMINPF_09287** | Transcriptional regulatory protein OmpR |
| **KCPMINPF_09336** | Transcriptional regulator MraZ |
| **KCPMINPF_09794** | Regulatory protein RecX |
| **KCPMINPF_09875** | LexA repressor |
| **KCPMINPF_09896** | Heat-inducible transcription repressor HrcA |
| **KCPMINPF_09913** | Transcriptional regulatory protein CusR |
| **KCPMINPF_100034** | Nitrogen regulatory protein P-II |
| **KCPMINPF_100042** | Transcriptional regulatory protein DegU |
| **KCPMINPF_10006** | Transcriptional repressor NrdR |
| **KCPMINPF_100095** | Peroxide-responsive repressor PerR |
| **KCPMINPF_100115** | Oxygen regulatory protein NreC |
| **KCPMINPF_10023** | Transcriptional regulator MraZ |
| **KCPMINPF_100316** | Transcriptional regulatory protein LnrK |
| **KCPMINPF_100329** | Transcriptional regulatory protein CusR |
| **KCPMINPF_100353** | RNA polymerase-binding transcription factor DksA |
| **KCPMINPF_100610** | Heat-inducible transcription repressor HrcA |
| **KCPMINPF_100669** | Regulatory protein MsrR |
| **KCPMINPF_100805** | Transcriptional activator protein CzcR |
| **KCPMINPF_100910** | RNA polymerase-binding transcription factor DksA |
| **KCPMINPF_100923** | Transcriptional regulatory protein LiaR |
| **KCPMINPF_101018** | Transcriptional regulatory protein WalR |
| **KCPMINPF_101141** | DnaA regulatory inactivator Hda |
| **KCPMINPF_101153** | Transcriptional regulatory protein WalR |
| **KCPMINPF_101238** | Transcriptional activator protein CopR |
| **KCPMINPF_101305** | Transcriptional repressor MprA |
| **KCPMINPF_101364** | Mercuric resistance operon regulatory protein |
| **KCPMINPF_101377** | LexA repressor |
| **KCPMINPF_101378** | LexA repressor |
| **KCPMINPF_101479** | Transcriptional regulatory protein LiaR |
| **KCPMINPF_101497** | RNA polymerase-binding transcription factor DksA |
| **KCPMINPF_101677** | Heat-inducible transcription repressor HrcA |
| **KCPMINPF_101709** | Alkaline phosphatase synthesis transcriptional regulatory protein PhoP |

| | |
|---|---|
| **KCPMINPF_101727** | Transcriptional regulatory protein DegU |
| **KCPMINPF_101778** | Transcriptional regulator SlyA |
| **KCPMINPF_101802** | Oxygen regulatory protein NreC |
| **KCPMINPF_101862** | Regulatory protein AtoC |
| **KCPMINPF_101870** | Bifunctional ligase/repressor BirA |
| **KCPMINPF_101933** | Transcriptional repressor NrdR |
| **KCPMINPF_102016** | Transcriptional regulatory protein KdpE |
| **KCPMINPF_102034** | Transcriptional regulatory protein LiaR |
| **KCPMINPF_102065** | Alkaline phosphatase synthesis transcriptional regulatory protein SphR |
| **KCPMINPF_10207** | Transcriptional regulatory protein LiaR |
| **KCPMINPF_102099** | ATP phosphoribosyltransferase regulatory subunit |
| **KCPMINPF_102305** | Iron-dependent repressor IdeR |
| **KCPMINPF_102379** | Alkaline phosphatase synthesis transcriptional regulatory protein PhoP |
| **KCPMINPF_102454** | Transcriptional regulatory protein tctD |
| **KCPMINPF_102468** | Transcriptional regulatory protein RcsB |
| **KCPMINPF_102654** | RNA polymerase-binding transcription factor CarD |
| **KCPMINPF_102789** | Glucitol operon repressor |
| **KCPMINPF_102849** | N-acetylglucosamine repressor |
| **KCPMINPF_102891** | Transcriptional regulatory protein SrrA |
| **KCPMINPF_102904** | Transcriptional regulatory protein DegU |
| **KCPMINPF_102960** | Transcriptional regulatory protein LiaR |
| **KCPMINPF_103119** | Transcriptional regulatory protein LiaR |
| **KCPMINPF_103161** | LexA repressor |
| **KCPMINPF_10320** | CdaA regulatory protein CdaR |
| **KCPMINPF_103220** | Transcriptional regulatory protein SrrA |
| **KCPMINPF_103306** | Regulatory protein RecX |
| **KCPMINPF_103311** | Heat-inducible transcription repressor HrcA |
| **KCPMINPF_103320** | Alkaline phosphatase synthesis transcriptional regulatory protein SphR |
| **KCPMINPF_103402** | Sensory/regulatory protein RpfC |
| **KCPMINPF_103414** | Transcriptional regulatory protein PhoP |
| **KCPMINPF_103575** | Transcriptional regulatory protein OmpR |
| **KCPMINPF_103583** | Penicillin-binding protein activator LpoA |
| **KCPMINPF_103609** | Transcriptional regulatory protein ZraR |
| **KCPMINPF_103616** | Transcriptional activator protein CopR |
| **KCPMINPF_10392** | Bifunctional ligase/repressor BirA |
| **KCPMINPF_103943** | Bifunctional transcriptional activator/DNA repair enzyme Ada |
| **KCPMINPF_103947** | Heat-inducible transcription repressor HrcA |
| **KCPMINPF_103960** | Glucitol operon repressor |
| **KCPMINPF_104117** | Mercuric resistance operon regulatory protein |
| **KCPMINPF_104159** | Transcriptional regulatory protein DegU |
| **KCPMINPF_104242** | RNA polymerase-binding transcription factor CarD |
| **KCPMINPF_104270** | Alkaline phosphatase synthesis transcriptional regulatory protein PhoP |
| **KCPMINPF_104298** | Redox-sensing transcriptional repressor Rex 1 |

| | |
|---|---|
| **KCPMINPF_104326** | Murein hydrolase activator EnvC |
| **KCPMINPF_104435** | Alkaline phosphatase synthesis transcriptional regulatory protein PhoP |
| **KCPMINPF_104462** | Bifunctional transcriptional activator/DNA repair enzyme Ada |
| **KCPMINPF_104487** | Transcriptional regulatory protein LiaR |
| **KCPMINPF_104552** | Transcriptional regulatory protein KdpE |
| **KCPMINPF_104569** | Transcriptional regulatory protein TcrA |
| **KCPMINPF_104665** | Transcriptional repressor SmtB |
| **KCPMINPF_104738** | Redox-sensing transcriptional repressor Rex 1 |
| **KCPMINPF_10476** | Murein hydrolase activator EnvC |
| **KCPMINPF_104981** | Nitrogen regulatory protein P-II |
| **KCPMINPF_105078** | KDP operon transcriptional regulatory protein KdpE |
| **KCPMINPF_105130** | Arginine repressor |
| **KCPMINPF_105269** | Transcriptional regulatory protein KdpE |
| **KCPMINPF_105271** | Transcriptional regulatory protein WalR |
| **KCPMINPF_105397** | Transcriptional regulatory protein DegU |
| **KCPMINPF_105477** | Anaerobic regulatory protein |
| **KCPMINPF_105524** | Transcriptional repressor NrdR |
| **KCPMINPF_105526** | LexA repressor |
| **KCPMINPF_105654** | Oxygen regulatory protein NreC |
| **KCPMINPF_105687** | Oxygen regulatory protein NreC |
| **KCPMINPF_105689** | Transcriptional regulatory protein LiaR |
| **KCPMINPF_105726** | Phosphate regulon transcriptional regulatory protein PhoB |
| **KCPMINPF_105761** | Transcriptional regulatory protein DegU |
| **KCPMINPF_105781** | Transcriptional regulatory protein DegU |
| **KCPMINPF_106058** | Heat-inducible transcription repressor HrcA |
| **KCPMINPF_106090** | Transcriptional regulatory protein KdpE |
| **KCPMINPF_106230** | Transcriptional regulatory protein LiaR |
| **KCPMINPF_106231** | Transcriptional regulatory protein LnrK |
| **KCPMINPF_106242** | Transcriptional regulatory protein LiaR |
| **KCPMINPF_106254** | Transcriptional repressor SmtB |
| **KCPMINPF_106318** | Transcriptional repressor SmtB |
| **KCPMINPF_106346** | Alkaline phosphatase synthesis transcriptional regulatory protein PhoP |
| **KCPMINPF_106570** | RNA polymerase-binding transcription factor DksA |
| **KCPMINPF_106683** | Transcriptional regulatory protein CusR |
| **KCPMINPF_106705** | Transcriptional regulatory protein DegU |
| **KCPMINPF_106737** | Transcriptional regulatory protein LiaR |
| **KCPMINPF_106899** | RNA polymerase-binding transcription factor CarD |
| **KCPMINPF_107106** | Transcriptional regulatory protein DegU |
| **KCPMINPF_107190** | Copper-sensing transcriptional repressor CsoR |
| **KCPMINPF_107208** | Alkaline phosphatase synthesis transcriptional regulatory protein PhoP |
| **KCPMINPF_107323** | Transcriptional regulatory protein HprR |
| **KCPMINPF_107451** | Photosynthetic apparatus regulatory protein RegA |
| **KCPMINPF_107488** | Ribose operon repressor |

| | |
|---|---|
| **KCPMINPF_107499** | Ribose operon repressor |
| **KCPMINPF_107571** | Transcriptional regulatory protein KdpE |
| **KCPMINPF_107640** | Regulatory protein AtoC |
| **KCPMINPF_107768** | Regulatory protein AtoC |
| **KCPMINPF_108102** | Alkaline phosphatase synthesis transcriptional regulatory protein PhoP |
| **KCPMINPF_108151** | LexA repressor |
| **KCPMINPF_108220** | LexA repressor |
| **KCPMINPF_108366** | Alkaline phosphatase synthesis transcriptional regulatory protein PhoP |
| **KCPMINPF_108450** | Alkaline phosphatase synthesis transcriptional regulatory protein SphR |
| **KCPMINPF_10847** | Transcriptional regulatory protein AfsQ1 |
| **KCPMINPF_10855** | Transcriptional regulator MraZ |
| **KCPMINPF_108655** | Transcriptional regulatory protein LiaR |
| **KCPMINPF_108708** | Transcriptional regulatory protein LnrK |
| **KCPMINPF_108769** | Transcriptional regulatory protein OmpR |
| **KCPMINPF_108884** | Redox-sensing transcriptional repressor Rex |
| **KCPMINPF_108939** | Transcriptional regulatory protein OmpR |
| **KCPMINPF_109002** | RNA polymerase-binding transcription factor DksA |
| **KCPMINPF_109053** | Transcriptional regulatory protein LiaR |
| **KCPMINPF_10930** | Alkaline phosphatase synthesis transcriptional regulatory protein PhoP |
| **KCPMINPF_109347** | Transcriptional regulatory protein DegU |
| **KCPMINPF_109374** | Bifunctional transcriptional activator/DNA repair enzyme Ada |
| **KCPMINPF_109620** | Transcriptional regulatory protein DesR |
| **KCPMINPF_109831** | Oxygen regulatory protein NreC |
| **KCPMINPF_10988** | Alkaline phosphatase synthesis transcriptional regulatory protein PhoP |
| **KCPMINPF_110015** | Oxygen regulatory protein NreC |
| **KCPMINPF_110035** | Transcriptional repressor PaaX |
| **KCPMINPF_110160** | DNA-binding transcriptional activator DevR/DosR |
| **KCPMINPF_110161** | Alkaline phosphatase synthesis transcriptional regulatory protein PhoP |
| **KCPMINPF_110195** | Transcriptional regulatory protein LiaR |
| **KCPMINPF_110206** | Transcriptional regulatory protein LiaR |
| **KCPMINPF_110415** | Transcriptional regulatory protein WalR |
| **KCPMINPF_110494** | Transcriptional regulatory protein WalR |
| **KCPMINPF_110495** | DNA-binding transcriptional activator DevR/DosR |
| **KCPMINPF_110502** | Oxygen regulatory protein NreC |
| **KCPMINPF_110589** | Penicillinase repressor |
| **KCPMINPF_110614** | Met repressor |
| **KCPMINPF_110785** | Transcriptional regulatory protein LiaR |
| **KCPMINPF_110823** | Mercuric resistance operon regulatory protein |
| **KCPMINPF_111024** | Lactose operon repressor |
| **KCPMINPF_111032** | Methanol dehydrogenase activator |
| **KCPMINPF_111040** | Redox-sensing transcriptional repressor Rex 1 |
| **KCPMINPF_111071** | DNA-binding transcriptional regulator NtrC |
| **KCPMINPF_111189** | Regulatory protein AtoC |

| | |
|---|---|
| **KCPMINPF_111507** | Phosphate regulon transcriptional regulatory protein PhoB |
| **KCPMINPF_111546** | Nitrogen regulatory protein P-II 1 |
| **KCPMINPF_111590** | Nitrogen regulatory protein |
| **KCPMINPF_111620** | Transcriptional activator HlyU |
| **KCPMINPF_11170** | Alkaline phosphatase synthesis transcriptional regulatory protein PhoP |
| **KCPMINPF_111719** | Oxygen regulatory protein NreC |
| **KCPMINPF_111761** | LexA repressor |
| **KCPMINPF_111788** | Transcriptional regulatory protein DegU |
| **KCPMINPF_111821** | Lactose operon repressor |
| **KCPMINPF_111944** | Alkaline phosphatase synthesis transcriptional regulatory protein PhoP |
| **KCPMINPF_111950** | Transcriptional regulatory protein PmpR |
| **KCPMINPF_112134** | Transcriptional regulatory protein FixJ |
| **KCPMINPF_112174** | Glucitol operon repressor |
| **KCPMINPF_112259** | Transcriptional regulatory protein LiaR |
| **KCPMINPF_11242** | Redox-sensing transcriptional repressor Rex |
| **KCPMINPF_112522** | Ribose operon repressor |
| **KCPMINPF_112596** | Transcriptional regulatory protein WalR |
| **KCPMINPF_112710** | Transcriptional regulatory protein DegU |
| **KCPMINPF_113414** | Transcriptional regulator KdgR |
| **KCPMINPF_11354** | ATP phosphoribosyltransferase regulatory subunit |
| **KCPMINPF_113549** | DNA-binding transcriptional activator DecR |
| **KCPMINPF_113563** | Oxygen regulatory protein NreC |
| **KCPMINPF_113604** | Ribose operon repressor |
| **KCPMINPF_113719** | Photosynthetic apparatus regulatory protein RegA |
| **KCPMINPF_113728** | Redox-sensing transcriptional repressor Rex 1 |
| **KCPMINPF_113817** | Alkaline phosphatase synthesis transcriptional regulatory protein PhoP |
| **KCPMINPF_113903** | Regulatory protein RecX |
| **KCPMINPF_114045** | Arabinose metabolism transcriptional repressor |
| **KCPMINPF_114059** | DNA-binding transcriptional activator DecR |
| **KCPMINPF_11409** | Phosphate regulon transcriptional regulatory protein PhoB |
| **KCPMINPF_114149** | Transcriptional regulatory protein WalR |
| **KCPMINPF_114166** | Transcriptional regulatory protein DegU |
| **KCPMINPF_114168** | Regulatory protein AtoC |
| **KCPMINPF_114246** | Transcriptional regulatory protein SrrA |
| **KCPMINPF_114288** | Transcriptional regulatory protein ZraR |
| **KCPMINPF_114301** | Iron-dependent repressor IdeR |
| **KCPMINPF_114346** | CdaA regulatory protein CdaR |
| **KCPMINPF_114474** | Transcriptional regulatory protein LiaR |
| **KCPMINPF_114527** | Mercuric resistance operon regulatory protein |
| **KCPMINPF_114636** | Oxygen regulatory protein NreC |
| **KCPMINPF_114639** | ATP phosphoribosyltransferase regulatory subunit |
| **KCPMINPF_114748** | Transcriptional regulator LsrR |
| **KCPMINPF_11475** | Transcriptional regulator MraZ |

| | |
|---|---|
| **KCPMINPF_114785** | Transcriptional activator protein CzcR |
| **KCPMINPF_114921** | Alkaline phosphatase synthesis transcriptional regulatory protein PhoP |
| **KCPMINPF_115006** | Transcriptional regulatory protein KdpE |
| **KCPMINPF_115101** | Transcriptional regulatory protein KdpE |
| **KCPMINPF_115108** | Transcriptional regulatory protein KdpE |
| **KCPMINPF_115229** | Hca operon transcriptional activator HcaR |
| **KCPMINPF_115412** | Oxygen regulatory protein NreC |
| **KCPMINPF_115453** | N-acetylglucosamine repressor |
| **KCPMINPF_115539** | Transcriptional regulatory protein KdpE |
| **KCPMINPF_115559** | Transcriptional regulatory protein LiaR |
| **KCPMINPF_115775** | Methanol dehydrogenase activator |
| **KCPMINPF_115788** | Transcriptional regulatory protein DegU |
| **KCPMINPF_115807** | Transcriptional regulatory protein LiaR |
| **KCPMINPF_115950** | Transcriptional regulatory protein SrrA |
| **KCPMINPF_116004** | Transcriptional regulatory protein WalR |
| **KCPMINPF_116084** | Bifunctional transcriptional activator/DNA repair enzyme Ada |
| **KCPMINPF_116514** | N-acetylglucosamine repressor |
| **KCPMINPF_116617** | Alkaline phosphatase synthesis transcriptional regulatory protein SphR |
| **KCPMINPF_116631** | Redox-sensing transcriptional repressor Rex |
| **KCPMINPF_116642** | Regulatory protein RecX |
| **KCPMINPF_116647** | Arabinose operon regulatory protein |
| **KCPMINPF_116656** | Transcriptional regulator MraZ |
| **KCPMINPF_116846** | Redox-sensing transcriptional repressor Rex 1 |
| **KCPMINPF_116879** | Transcriptional regulatory protein LnrK |
| **KCPMINPF_116950** | Transcriptional regulator SlyA |
| **KCPMINPF_11700** | LexA repressor |
| **KCPMINPF_117261** | Transcriptional repressor NrdR |
| **KCPMINPF_117602** | Transcriptional regulatory protein OmpR |
| **KCPMINPF_117837** | Transcriptional regulatory protein SrrA |
| **KCPMINPF_117889** | Transcriptional regulator WhiB |
| **KCPMINPF_117977** | Transcriptional regulatory protein LiaR |
| **KCPMINPF_118040** | Copper-sensing transcriptional repressor CsoR |
| **KCPMINPF_118147** | Transcriptional activator protein CopR |
| **KCPMINPF_118195** | Ribose operon repressor |
| **KCPMINPF_118329** | Pca regulon regulatory protein |
| **KCPMINPF_11833** | Transcriptional regulatory protein PmpR |
| **KCPMINPF_118357** | Oxygen regulatory protein NreC |
| **KCPMINPF_118514** | Transcriptional regulatory protein CseB |
| **KCPMINPF_118639** | Transcriptional regulatory protein LnrK |
| **KCPMINPF_118865** | Phosphate regulon transcriptional regulatory protein PhoB |
| **KCPMINPF_118895** | Glucitol operon repressor |
| **KCPMINPF_118955** | Alkaline phosphatase synthesis transcriptional regulatory protein PhoP |
| **KCPMINPF_119051** | Transcriptional regulatory protein WalR |

| | |
|---|---|
| **KCPMINPF_119154** | Alkaline phosphatase synthesis transcriptional regulatory protein PhoP |
| **KCPMINPF_119170** | Pca regulon regulatory protein |
| **KCPMINPF_119292** | Oxygen regulatory protein NreC |
| **KCPMINPF_119381** | Heat-inducible transcription repressor HrcA |
| **KCPMINPF_119499** | Transcriptional regulatory protein DegU |
| **KCPMINPF_119513** | Alkaline phosphatase synthesis transcriptional regulatory protein PhoP |
| **KCPMINPF_119586** | Oxygen regulatory protein NreC |
| **KCPMINPF_119628** | Nitrogen regulatory protein P-II |
| **KCPMINPF_119778** | RNA polymerase-binding transcription factor CarD |
| **KCPMINPF_119911** | Alkaline phosphatase synthesis transcriptional regulatory protein SphR |
| **KCPMINPF_120191** | Transcriptional regulatory protein DegU |
| **KCPMINPF_120206** | Oxygen regulatory protein NreC |
| **KCPMINPF_120404** | Luminescence regulatory protein LuxO |
| **KCPMINPF_120608** | Regulatory protein RecX |
| **KCPMINPF_120717** | Regulatory protein AtoC |
| **KCPMINPF_120847** | Nitrogen regulatory protein |
| **KCPMINPF_121039** | Transcriptional regulator KdgR |
| **KCPMINPF_121057** | Transcriptional regulatory protein LiaR |
| **KCPMINPF_121216** | Oxygen regulatory protein NreC |
| **KCPMINPF_121221** | Transcriptional regulatory protein LiaR |
| **KCPMINPF_12133** | Methanol dehydrogenase activator |
| **KCPMINPF_121435** | Phosphoenolpyruvate synthase regulatory protein |
| **KCPMINPF_121528** | Regulatory protein RecX |
| **KCPMINPF_121692** | Transcriptional activator protein CopR |
| **KCPMINPF_121769** | Regulatory protein RecX |
| **KCPMINPF_12193** | Regulatory protein AtoC |
| **KCPMINPF_121943** | Transcriptional regulatory protein LiaR |
| **KCPMINPF_121994** | Transcriptional regulatory protein LiaR |
| **KCPMINPF_122008** | Transcriptional regulatory protein WalR |
| **KCPMINPF_12210** | Regulatory protein AtoC |
| **KCPMINPF_122260** | Heat-inducible transcription repressor HrcA |
| **KCPMINPF_122403** | Oxygen regulatory protein NreC |
| **KCPMINPF_122591** | Nitrogen regulatory protein P-II 2 |
| **KCPMINPF_122607** | Redox-sensing transcriptional repressor Rex 1 |
| **KCPMINPF_122739** | Regulatory protein AtoC |
| **KCPMINPF_122778** | Transcriptional repressor NrdR |
| **KCPMINPF_122819** | N-acetylglucosamine repressor |
| **KCPMINPF_122942** | Mercuric resistance operon regulatory protein |
| **KCPMINPF_122959** | Leucine-responsive regulatory protein |
| **KCPMINPF_123215** | Alkaline phosphatase synthesis transcriptional regulatory protein PhoP |
| **KCPMINPF_123255** | Transcriptional regulatory protein QseB |
| **KCPMINPF_123322** | Transcriptional regulatory protein LiaR |
| **KCPMINPF_123457** | Copper-sensing transcriptional repressor CsoR |

| | |
|---|---|
| **KCPMINPF_123484** | Transcriptional regulatory protein ros |
| **KCPMINPF_123501** | Alkaline phosphatase synthesis transcriptional regulatory protein PhoP |
| **KCPMINPF_123573** | Transcriptional regulatory protein WalR |
| **KCPMINPF_12359** | Alkaline phosphatase synthesis transcriptional regulatory protein PhoP |
| **KCPMINPF_123664** | KDP operon transcriptional regulatory protein KdpE |
| **KCPMINPF_123738** | Alkaline phosphatase synthesis transcriptional regulatory protein PhoP |
| **KCPMINPF_123751** | Transcriptional regulatory protein LiaR |
| **KCPMINPF_12437** | Alkaline phosphatase synthesis transcriptional regulatory protein PhoP |
| **KCPMINPF_124548** | Transcriptional regulatory protein LiaR |
| **KCPMINPF_124609** | Negative regulatory protein YxlE |
| **KCPMINPF_124668** | Transcriptional regulatory protein BaeR |
| **KCPMINPF_124695** | Alkaline phosphatase synthesis transcriptional regulatory protein PhoP |
| **KCPMINPF_12480** | Transcriptional repressor SmtB |
| **KCPMINPF_124866** | Transcriptional regulatory protein ZraR |
| **KCPMINPF_124969** | Transcriptional regulatory protein OmpR |
| **KCPMINPF_12504** | Peroxide-responsive repressor PerR |
| **KCPMINPF_125113** | Leucine-responsive regulatory protein |
| **KCPMINPF_125287** | Lactose operon repressor |
| **KCPMINPF_125348** | Lactose operon repressor |
| **KCPMINPF_125443** | Transcriptional regulatory protein TcrA |
| **KCPMINPF_125480** | Leucine-responsive regulatory protein |
| **KCPMINPF_12549** | Transcriptional activator protein CzcR |
| **KCPMINPF_125505** | Transcriptional regulatory protein WalR |
| **KCPMINPF_125509** | Transcriptional regulatory protein DegU |
| **KCPMINPF_12562** | Heat-inducible transcription repressor HrcA |
| **KCPMINPF_125639** | LexA repressor |
| **KCPMINPF_12569** | Pur operon repressor |
| **KCPMINPF_125704** | Transcriptional regulatory protein LnrK |
| **KCPMINPF_125716** | Hydrogen peroxide-inducible genes activator |
| **KCPMINPF_125723** | Oxygen regulatory protein NreC |
| **KCPMINPF_125822** | Bifunctional ligase/repressor BirA |
| **KCPMINPF_125902** | Transcriptional regulatory protein QseB |
| **KCPMINPF_125919** | Transcriptional regulatory protein LiaR |
| **KCPMINPF_125931** | Alkaline phosphatase synthesis transcriptional regulatory protein PhoP |
| **KCPMINPF_12600** | PTS-dependent dihydroxyacetone kinase operon regulatory protein |
| **KCPMINPF_126061** | Transcriptional regulatory protein DegU |
| **KCPMINPF_126544** | Oxygen regulatory protein NreC |
| **KCPMINPF_126575** | Alkaline phosphatase synthesis transcriptional regulatory protein SphR |
| **KCPMINPF_126703** | DnaA regulatory inactivator Hda |
| **KCPMINPF_126714** | Negative regulatory protein YxlE |
| **KCPMINPF_12685** | Biofilm growth-associated repressor |
| **KCPMINPF_126913** | Murein hydrolase activator NlpD |
| **KCPMINPF_12692** | Oxygen regulatory protein NreC |

| | |
|---|---|
| **KCPMINPF_126938** | Alkaline phosphatase synthesis transcriptional regulatory protein PhoP |
| **KCPMINPF_12703** | Transcriptional regulatory protein DegU |
| **KCPMINPF_127153** | Virulence transcriptional regulatory protein PhoP |
| **KCPMINPF_127157** | LexA repressor |
| **KCPMINPF_127326** | RNA polymerase-binding transcription factor CarD |
| **KCPMINPF_127452** | Sigma54-dependent transcriptional activator SfnR |
| **KCPMINPF_12751** | Transcriptional regulatory protein WalR |
| **KCPMINPF_127551** | Alkaline phosphatase synthesis transcriptional regulatory protein PhoP |
| **KCPMINPF_127577** | Copper-sensing transcriptional repressor CsoR |
| **KCPMINPF_12764** | Regulatory protein RecX |
| **KCPMINPF_12769** | Transcriptional regulatory protein WalR |
| **KCPMINPF_127693** | Transcriptional regulatory protein WalR |
| **KCPMINPF_127695** | Regulatory protein AtoC |
| **KCPMINPF_127704** | Tetrathionate response regulatory protein TtrR |
| **KCPMINPF_127705** | Transcriptional regulatory protein TdiR |
| **KCPMINPF_127719** | Transcriptional activator NphR |
| **KCPMINPF_127749** | KDP operon transcriptional regulatory protein KdpE |
| **KCPMINPF_127815** | Transcriptional regulatory protein DegU |
| **KCPMINPF_127849** | Organic hydroperoxide resistance transcriptional regulator |
| **KCPMINPF_128038** | Transcriptional regulatory protein BaeR |
| **KCPMINPF_128112** | DNA-binding transcriptional regulator NtrC |
| **KCPMINPF_128143** | Transcriptional repressor NrdR |
| **KCPMINPF_128215** | Regulatory protein AtoC |
| **KCPMINPF_128269** | Transcriptional regulatory protein LiaR |
| **KCPMINPF_12827** | Transcriptional regulatory protein DegU |
| **KCPMINPF_128398** | Alkaline phosphatase synthesis transcriptional regulatory protein PhoP |
| **KCPMINPF_128508** | DNA-binding transcriptional activator DecR |
| **KCPMINPF_12851** | Transcriptional regulatory protein LnrK |
| **KCPMINPF_128534** | Alkaline phosphatase synthesis transcriptional regulatory protein PhoP |
| **KCPMINPF_128583** | Transcriptional regulator MraZ |
| **KCPMINPF_128812** | Transcriptional regulatory protein WalR |
| **KCPMINPF_128822** | (R)-phenyllactate dehydratase activator |
| **KCPMINPF_128823** | (R)-phenyllactate dehydratase activator |
| **KCPMINPF_128828** | Nitrogen regulatory protein P-II 2 |
| **KCPMINPF_128901** | Alkaline phosphatase synthesis transcriptional regulatory protein PhoP |
| **KCPMINPF_128958** | Regulatory protein RecX |
| **KCPMINPF_129107** | Transcriptional regulatory protein ZraR |
| **KCPMINPF_129181** | Biofilm growth-associated repressor |
| **KCPMINPF_129283** | Transcriptional regulator LsrR |
| **KCPMINPF_129352** | Transcriptional repressor SmtB |
| **KCPMINPF_129576** | KDP operon transcriptional regulatory protein KdpE |
| **KCPMINPF_129684** | Sensory/regulatory protein RpfC |
| **KCPMINPF_129703** | Alkaline phosphatase synthesis transcriptional regulatory protein PhoP |

| | |
|---|---|
| **KCPMINPF_129932** | Transcriptional regulatory protein LiaR |
| **KCPMINPF_130040** | Lactose operon repressor |
| **KCPMINPF_130063** | Hydrogen peroxide-inducible genes activator |
| **KCPMINPF_130107** | Transcriptional regulatory protein DegU |
| **KCPMINPF_130409** | Heat-inducible transcription repressor HrcA |
| **KCPMINPF_130485** | Purine catabolism regulatory protein |
| **KCPMINPF_130501** | Transcriptional activator HlyU |
| **KCPMINPF_130542** | Transcriptional regulatory protein BtsR |
| **KCPMINPF_130547** | Bifunctional transcriptional activator/DNA repair enzyme Ada |
| **KCPMINPF_131121** | Transcriptional regulatory protein DegU |
| **KCPMINPF_131188** | Transcriptional regulatory protein DegU |
| **KCPMINPF_131403** | N-acetylglucosamine repressor |
| **KCPMINPF_131637** | Regulatory protein Spx |
| **KCPMINPF_131749** | Transcriptional activator protein CopR |
| **KCPMINPF_131818** | Nitrogen regulatory protein P-II |
| **KCPMINPF_131845** | LexA repressor |
| **KCPMINPF_132017** | Gliding motility regulatory protein |
| **KCPMINPF_132026** | Transcriptional regulator BlaI |
| **KCPMINPF_132028** | LexA repressor |
| **KCPMINPF_132283** | LexA repressor |
| **KCPMINPF_13232** | Transcriptional regulatory protein DegU |
| **KCPMINPF_132332** | Bifunctional ligase/repressor BirA |
| **KCPMINPF_132515** | LexA repressor |
| **KCPMINPF_132581** | Glycine cleavage system transcriptional activator |
| **KCPMINPF_13262** | Murein hydrolase activator EnvC |
| **KCPMINPF_132657** | Pyruvate dehydrogenase complex repressor |
| **KCPMINPF_132848** | Transcriptional regulator AcuR |
| **KCPMINPF_132880** | DNA-binding transcriptional regulator BolA |
| **KCPMINPF_132895** | Transcriptional regulatory protein KdpE |
| **KCPMINPF_132914** | Transcriptional regulatory protein WalR |
| **KCPMINPF_132964** | Glycine cleavage system transcriptional activator |
| **KCPMINPF_133106** | DNA-binding transcriptional regulator BolA |
| **KCPMINPF_13319** | Leucine-responsive regulatory protein |
| **KCPMINPF_133232** | Transcriptional activator HlyU |
| **KCPMINPF_133800** | Transcriptional regulatory protein LiaR |
| **KCPMINPF_133815** | Transcriptional regulatory protein LiaR |
| **KCPMINPF_13383** | Transcriptional regulatory protein ros |
| **KCPMINPF_133846** | Transcriptional regulatory protein ZraR |
| **KCPMINPF_133923** | RNA polymerase-binding transcription factor CarD |
| **KCPMINPF_133925** | Transcriptional regulatory protein ZraR |
| **KCPMINPF_13405** | Transcriptional regulatory protein OmpR |
| **KCPMINPF_134149** | RNA polymerase-binding transcription factor DksA |
| **KCPMINPF_134226** | Phosphate regulon transcriptional regulatory protein PhoB |

| | |
|---|---|
| **KCPMINPF_134257** | N-acetylglucosamine repressor |
| **KCPMINPF_134284** | N-acetylglucosamine repressor |
| **KCPMINPF_13453** | DNA-binding transcriptional regulator NtrC |
| **KCPMINPF_13455** | DNA-binding transcriptional regulator NtrC |
| **KCPMINPF_13480** | LexA repressor |
| **KCPMINPF_134831** | RNA polymerase-binding transcription factor CarD |
| **KCPMINPF_134839** | Alkaline phosphatase synthesis transcriptional regulatory protein SphR |
| **KCPMINPF_135134** | Nitrogen regulatory protein P-II |
| **KCPMINPF_135145** | Ribose operon repressor |
| **KCPMINPF_135163** | Transcriptional regulatory protein BtsR |
| **KCPMINPF_13523** | Transcriptional regulatory protein AfsQ1 |
| **KCPMINPF_135283** | Transcriptional regulatory protein CusR |
| **KCPMINPF_135308** | Transcriptional regulatory protein LiaR |
| **KCPMINPF_135424** | N-acetylglucosamine repressor |
| **KCPMINPF_135501** | Transcriptional regulatory protein DesR |
| **KCPMINPF_135513** | Transcriptional regulator SlyA |
| **KCPMINPF_135554** | Transcriptional repressor IclR |
| **KCPMINPF_135560** | Sensory/regulatory protein RpfC |
| **KCPMINPF_135697** | N-acetylglucosamine repressor |
| **KCPMINPF_135737** | N-acetylglucosamine repressor |
| **KCPMINPF_13597** | Bifunctional ligase/repressor BirA |
| **KCPMINPF_135982** | Ribose operon repressor |
| **KCPMINPF_136083** | Oxygen regulatory protein NreC |
| **KCPMINPF_136128** | Transcriptional regulatory protein WalR |
| **KCPMINPF_136192** | Regulatory protein AtoC |
| **KCPMINPF_136265** | Transcriptional regulatory protein BtsR |
| **KCPMINPF_136286** | Transcriptional regulatory protein WalR |
| **KCPMINPF_13633** | Nitrogen regulatory protein P-II |
| **KCPMINPF_136389** | Transcriptional activatory protein BadR |
| **KCPMINPF_136562** | Transcriptional regulatory protein WalR |
| **KCPMINPF_136642** | RNA polymerase-binding transcription factor DksA |
| **KCPMINPF_136658** | Transcriptional regulator KdgR |
| **KCPMINPF_136714** | RNA polymerase-binding transcription factor DksA |
| **KCPMINPF_136860** | Bifunctional ligase/repressor BirA |
| **KCPMINPF_136869** | Psp operon transcriptional activator |
| **KCPMINPF_136973** | Transcriptional regulatory protein DesR |
| **KCPMINPF_136976** | Transcriptional repressor NrdR |
| **KCPMINPF_137078** | Transcriptional regulatory protein PrrA |
| **KCPMINPF_137144** | Transcriptional repressor SmtB |
| **KCPMINPF_137243** | Methanol dehydrogenase activator |
| **KCPMINPF_13727** | RNA polymerase-binding transcription factor DksA |
| **KCPMINPF_137271** | Transcriptional regulatory protein LnrK |
| **KCPMINPF_137304** | Transcriptional regulatory protein DegU |

| | |
|---|---|
| **KCPMINPF_137305** | Transcriptional regulatory protein DegU |
| **KCPMINPF_13743** | Regulatory protein AtoC |
| **KCPMINPF_13745** | Transcriptional regulatory protein AfsQ1 |
| **KCPMINPF_137549** | Peroxide-responsive repressor PerR |
| **KCPMINPF_137688** | DNA-binding transcriptional regulator BolA |
| **KCPMINPF_137894** | Transcriptional regulatory protein QseB |
| **KCPMINPF_137930** | Phosphate regulon transcriptional regulatory protein PhoB |
| **KCPMINPF_138306** | Phosphate regulon transcriptional regulatory protein PhoB |
| **KCPMINPF_138547** | Bifunctional transcriptional activator/DNA repair enzyme Ada |
| **KCPMINPF_138564** | Glycerol-3-phosphate regulon repressor |
| **KCPMINPF_138598** | Transcriptional activator protein CzcR |
| **KCPMINPF_138636** | LexA repressor |
| **KCPMINPF_138749** | Transcriptional regulatory protein OmpR |
| **KCPMINPF_139117** | Gliding motility regulatory protein |
| **KCPMINPF_139149** | Transcriptional repressor NrdR |
| **KCPMINPF_139211** | Transcriptional regulatory protein WalR |
| **KCPMINPF_139235** | Alkaline phosphatase synthesis transcriptional regulatory protein SphR |
| **KCPMINPF_139246** | Nitrogen regulatory protein P-II |
| **KCPMINPF_139287** | Lactose operon repressor |
| **KCPMINPF_139367** | Transcriptional repressor FrmR |
| **KCPMINPF_139515** | Transcriptional regulatory protein KdpE |
| **KCPMINPF_139725** | Transcriptional regulatory protein BaeR |
| **KCPMINPF_139990** | Transcriptional regulatory protein DegU |
| **KCPMINPF_140269** | Pca regulon regulatory protein |
| **KCPMINPF_140489** | Alkaline phosphatase synthesis transcriptional regulatory protein PhoP |
| **KCPMINPF_140560** | Alkaline phosphatase synthesis transcriptional regulatory protein PhoP |
| **KCPMINPF_140567** | Glc operon transcriptional activator |
| **KCPMINPF_140687** | Glycerol-3-phosphate regulon repressor |
| **KCPMINPF_14079** | LexA repressor |
| **KCPMINPF_141147** | Transcriptional regulatory protein BaeR |
| **KCPMINPF_14116** | Regulatory protein AtoC |
| **KCPMINPF_141281** | Transcriptional activatory protein AadR |
| **KCPMINPF_14134** | LexA repressor |
| **KCPMINPF_141399** | Transcriptional regulatory protein ros |
| **KCPMINPF_141449** | Transcriptional regulatory protein WalR |
| **KCPMINPF_141698** | Transcriptional repressor PaaX |
| **KCPMINPF_14176** | Phosphate regulon transcriptional regulatory protein PhoB |
| **KCPMINPF_141857** | Phosphoglycerate transport regulatory protein PgtC |
| **KCPMINPF_141913** | Transcriptional regulatory protein FixJ |
| **KCPMINPF_142038** | Nitrogen regulatory protein P-II |
| **KCPMINPF_142106** | Heat-inducible transcription repressor HrcA |
| **KCPMINPF_142257** | Sensory/regulatory protein RpfC |
| **KCPMINPF_142350** | Transcriptional regulatory protein WalR |

| | |
|---|---|
| **KCPMINPF_142393** | Transcriptional activator protein CopR |
| **KCPMINPF_142440** | Regulatory protein RecX |
| **KCPMINPF_142567** | Transcriptional regulatory protein DegU |
| **KCPMINPF_142654** | Copper-sensing transcriptional repressor CsoR |
| **KCPMINPF_142789** | Tetracycline repressor protein class G |
| **KCPMINPF_142884** | Nitrogen regulatory protein P-II 2 |
| **KCPMINPF_14294** | Transcriptional regulatory protein ZraR |
| **KCPMINPF_143054** | Regulatory protein AtoC |
| **KCPMINPF_143491** | Heat-inducible transcription repressor HrcA |
| **KCPMINPF_143512** | DNA-binding transcriptional regulator NtrC |
| **KCPMINPF_143577** | Iron-dependent repressor IdeR |
| **KCPMINPF_143595** | Glucitol operon repressor |
| **KCPMINPF_143615** | Penicillinase repressor |
| **KCPMINPF_143700** | Transcriptional regulatory protein DegU |
| **KCPMINPF_143802** | Heat-inducible transcription repressor HrcA |
| **KCPMINPF_143830** | Oxygen regulatory protein NreC |
| **KCPMINPF_143868** | Oxygen regulatory protein NreC |
| **KCPMINPF_144241** | Transcriptional regulatory protein WalR |
| **KCPMINPF_144359** | Sigma factor AlgU regulatory protein MucB |
| **KCPMINPF_144369** | Extracellular matrix regulatory protein A |
| **KCPMINPF_144413** | Transcriptional regulator KdgR |
| **KCPMINPF_144427** | ATP phosphoribosyltransferase regulatory subunit |
| **KCPMINPF_144473** | Bifunctional transcriptional activator/DNA repair enzyme Ada |
| **KCPMINPF_144595** | Peroxide-responsive repressor PerR |
| **KCPMINPF_144598** | Peroxide-responsive repressor PerR |
| **KCPMINPF_14481** | Methanol dehydrogenase activator |
| **KCPMINPF_144829** | Transcriptional repressor NrdR |
| **KCPMINPF_145066** | Transcriptional regulator MraZ |
| **KCPMINPF_145144** | Mercuric resistance operon regulatory protein |
| **KCPMINPF_14551** | LexA repressor |
| **KCPMINPF_145550** | Nitrogen regulatory protein P-II 1 |
| **KCPMINPF_145586** | Hydrogen peroxide-inducible genes activator |
| **KCPMINPF_14562** | Regulatory protein RecX |
| **KCPMINPF_145750** | Transcriptional regulatory protein KdpE |
| **KCPMINPF_145808** | N-acetylglucosamine repressor |
| **KCPMINPF_146099** | Glc operon transcriptional activator |
| **KCPMINPF_146184** | Transcriptional regulatory protein WalR |
| **KCPMINPF_146314** | Glycine cleavage system transcriptional activator |
| **KCPMINPF_14635** | Copper-sensing transcriptional repressor CsoR |
| **KCPMINPF_146468** | Transcriptional regulatory protein OmpR |
| **KCPMINPF_146557** | LexA repressor |
| **KCPMINPF_146618** | Bifunctional ligase/repressor BirA |
| **KCPMINPF_146664** | Transcriptional regulatory protein WalR |

| | |
|---|---|
| **KCPMINPF_146842** | Hydrogen peroxide-inducible genes activator |
| **KCPMINPF_146955** | Iron-dependent repressor IdeR |
| **KCPMINPF_146956** | Iron-dependent repressor IdeR |
| **KCPMINPF_147142** | Hca operon transcriptional activator HcaR |
| **KCPMINPF_147332** | RNA polymerase-binding transcription factor DksA |
| **KCPMINPF_147445** | Transcriptional repressor SmtB |
| **KCPMINPF_147542** | Transcriptional regulatory protein OmpR |
| **KCPMINPF_14755** | Transcriptional regulatory protein WalR |
| **KCPMINPF_147596** | Transcriptional repressor NrdR |
| **KCPMINPF_147643** | Alkaline phosphatase synthesis transcriptional regulatory protein SphR |
| **KCPMINPF_147917** | Transcriptional regulatory protein KdpE |
| **KCPMINPF_148065** | CdaA regulatory protein CdaR |
| **KCPMINPF_148087** | Denitrification regulatory protein NirQ |
| **KCPMINPF_148188** | Oxygen regulatory protein NreC |
| **KCPMINPF_148630** | Ribose operon repressor |
| **KCPMINPF_148695** | Glycerol-3-phosphate regulon repressor |
| **KCPMINPF_148917** | Transcriptional repressor SmtB |
| **KCPMINPF_14896** | Transcriptional regulatory protein ZraR |
| **KCPMINPF_149005** | Transcriptional regulator SlyA |
| **KCPMINPF_149006** | Transcriptional regulator SlyA |
| **KCPMINPF_149057** | Transcriptional activator protein CopR |
| **KCPMINPF_149204** | Transcriptional regulatory protein WalR |
| **KCPMINPF_149398** | Transcriptional regulatory protein WalR |
| **KCPMINPF_14943** | Transcriptional regulatory protein BtsR |
| **KCPMINPF_14947** | Regulatory protein AtoC |
| **KCPMINPF_149514** | Alkaline phosphatase synthesis transcriptional regulatory protein PhoP |
| **KCPMINPF_149655** | Alkaline phosphatase synthesis transcriptional regulatory protein PhoP |
| **KCPMINPF_149735** | Oxygen regulatory protein NreC |
| **KCPMINPF_149777** | Transcriptional regulatory protein LnrK |
| **KCPMINPF_149901** | Transcriptional regulatory protein DegU |
| **KCPMINPF_15028** | Denitrification regulatory protein NirQ |
| **KCPMINPF_150351** | Leucine-responsive regulatory protein |
| **KCPMINPF_150389** | cAMP-activated global transcriptional regulator CRP |
| **KCPMINPF_150558** | Transcriptional repressor NrdR |
| **KCPMINPF_15065** | Hydrogen peroxide-inducible genes activator |
| **KCPMINPF_150753** | Bifunctional ligase/repressor BirA |
| **KCPMINPF_150894** | Transcriptional regulatory protein PhoP |
| **KCPMINPF_15128** | Transcriptional regulatory protein YpdB |
| **KCPMINPF_151416** | LexA repressor |
| **KCPMINPF_151463** | KDP operon transcriptional regulatory protein KdpE |
| **KCPMINPF_151511** | Transcriptional regulatory protein HprR |
| **KCPMINPF_151634** | Bifunctional ligase/repressor BirA |
| **KCPMINPF_151665** | Alkaline phosphatase synthesis transcriptional regulatory protein PhoP |

| | |
|---|---|
| **KCPMINPF_15171** | Murein hydrolase activator EnvC |
| **KCPMINPF_151748** | Exoenzyme S synthesis regulatory protein ExsA |
| **KCPMINPF_151779** | Transcriptional regulatory protein DegU |
| **KCPMINPF_151856** | Transcriptional regulatory protein FixJ |
| **KCPMINPF_151857** | Transcriptional regulatory protein TdiR |
| **KCPMINPF_151931** | Heat-inducible transcription repressor HrcA |
| **KCPMINPF_152113** | Sensory/regulatory protein RpfC |
| **KCPMINPF_15215** | Phosphate regulon transcriptional regulatory protein PhoB |
| **KCPMINPF_15219** | Transcriptional regulatory protein LiaR |
| **KCPMINPF_152919** | Transcriptional regulator MraZ |
| **KCPMINPF_152979** | Oxygen regulatory protein NreC |
| **KCPMINPF_15309** | Transcriptional regulatory protein KdpE |
| **KCPMINPF_153251** | Nitrogen regulatory protein P-II 2 |
| **KCPMINPF_153398** | Bifunctional ligase/repressor BirA |
| **KCPMINPF_153675** | ATP phosphoribosyltransferase regulatory subunit |
| **KCPMINPF_153707** | Heat-inducible transcription repressor HrcA |
| **KCPMINPF_15431** | Transcriptional regulatory protein ZraR |
| **KCPMINPF_154332** | Oxygen regulatory protein NreC |
| **KCPMINPF_154384** | Copper-sensing transcriptional repressor CsoR |
| **KCPMINPF_154452** | Transcriptional regulatory protein FixJ |
| **KCPMINPF_15449** | Transcriptional regulatory protein WalR |
| **KCPMINPF_154635** | Formate hydrogenlyase transcriptional activator FhlA |
| **KCPMINPF_154810** | Transcriptional regulator SlyA |
| **KCPMINPF_15485** | Bifunctional ligase/repressor BirA |
| **KCPMINPF_154850** | Glucitol operon repressor |
| **KCPMINPF_154859** | Erythritol catabolism regulatory protein EryD |
| **KCPMINPF_155161** | Transcriptional regulatory protein BtsR |
| **KCPMINPF_15551** | Transcriptional regulatory protein SrrA |
| **KCPMINPF_155545** | Alkaline phosphatase synthesis transcriptional regulatory protein PhoP |
| **KCPMINPF_155589** | Transcriptional activator protein NhaR |
| **KCPMINPF_155637** | Iron-dependent repressor IdeR |
| **KCPMINPF_155794** | Transcriptional regulatory protein LiaR |
| **KCPMINPF_155797** | Transcriptional regulatory protein SrrA |
| **KCPMINPF_155933** | Sensory/regulatory protein RpfC |
| **KCPMINPF_155984** | Alkaline phosphatase synthesis transcriptional regulatory protein PhoP |
| **KCPMINPF_156098** | Heat-inducible transcription repressor HrcA |
| **KCPMINPF_156103** | Glycine cleavage system transcriptional activator |
| **KCPMINPF_15618** | LexA repressor |
| **KCPMINPF_156254** | Transcriptional regulatory protein QseB |
| **KCPMINPF_156270** | N-acetylglucosamine repressor |
| **KCPMINPF_15649** | Regulatory protein AtoC |
| **KCPMINPF_15652** | Transcriptional regulatory protein BtsR |
| **KCPMINPF_156563** | RNA polymerase-binding transcription factor DksA |

| | |
|---|---|
| **KCPMINPF_156752** | Oxygen regulatory protein NreC |
| **KCPMINPF_156768** | Transcriptional regulatory protein FixJ |
| **KCPMINPF_156902** | Transcriptional regulatory protein KdpE |
| **KCPMINPF_156955** | LexA repressor |
| **KCPMINPF_157052** | Transcriptional regulatory protein WalR |
| **KCPMINPF_157069** | Transcriptional repressor NrdR |
| **KCPMINPF_157140** | Transcriptional regulator SlyA |
| **KCPMINPF_157189** | Bifunctional ligase/repressor BirA |
| **KCPMINPF_15741** | RNA polymerase-binding transcription factor DksA |
| **KCPMINPF_157417** | Transcriptional regulatory protein LiaR |
| **KCPMINPF_15744** | Bifunctional transcriptional activator/DNA repair enzyme Ada |
| **KCPMINPF_157513** | Oxygen regulatory protein NreC |
| **KCPMINPF_157530** | DNA-binding transcriptional regulator NtrC |
| **KCPMINPF_15785** | Transcriptional regulatory protein ZraR |
| **KCPMINPF_157913** | Transcriptional regulator MraZ |
| **KCPMINPF_157957** | Transcriptional regulator MntR |
| **KCPMINPF_158082** | LexA repressor |
| **KCPMINPF_158109** | Nitrogen regulatory protein |
| **KCPMINPF_158183** | LexA repressor |
| **KCPMINPF_15838** | DNA-binding transcriptional regulator NtrC |
| **KCPMINPF_158479** | Pca regulon regulatory protein |
| **KCPMINPF_15884** | Transcriptional regulator MraZ |
| **KCPMINPF_158863** | Hydrogen peroxide-inducible genes activator |
| **KCPMINPF_159099** | LexA repressor |
| **KCPMINPF_159281** | Murein hydrolase activator NlpD |
| **KCPMINPF_159768** | Transcriptional regulatory protein YpdB |
| **KCPMINPF_160023** | Transcriptional regulatory protein QseB |
| **KCPMINPF_160131** | Copper-sensing transcriptional repressor CsoR |
| **KCPMINPF_16025** | Transcriptional regulator SlyA |
| **KCPMINPF_160321** | Transcriptional repressor NrdR |
| **KCPMINPF_160361** | Glucitol operon repressor |
| **KCPMINPF_160832** | Regulatory protein AtoC |
| **KCPMINPF_16084** | Transcriptional regulatory protein FixJ |
| **KCPMINPF_160857** | Transcriptional regulatory protein DegU |
| **KCPMINPF_160930** | Transcriptional regulatory protein DegU |
| **KCPMINPF_161389** | Anaerobic regulatory protein |
| **KCPMINPF_16144** | Regulatory protein AtoC |
| **KCPMINPF_161630** | Transcriptional regulatory protein QseB |
| **KCPMINPF_16166** | flagellum biosynthesis repressor protein FlbT |
| **KCPMINPF_16171** | Flagellar transcriptional regulator FtcR |
| **KCPMINPF_161755** | Oxygen regulatory protein NreC |
| **KCPMINPF_161834** | Oxygen regulatory protein NreC |
| **KCPMINPF_161885** | Oxygen regulatory protein NreC |

| | |
|---|---|
| **KCPMINPF_162049** | LexA repressor |
| **KCPMINPF_16214** | Transcriptional repressor SmtB |
| **KCPMINPF_162261** | Transcriptional regulatory protein DegU |
| **KCPMINPF_162332** | Alkaline phosphatase synthesis transcriptional regulatory protein PhoP |
| **KCPMINPF_162387** | Phosphate regulon transcriptional regulatory protein PhoB |
| **KCPMINPF_162389** | Transcriptional regulatory protein BtsR |
| **KCPMINPF_162579** | Alginate biosynthesis transcriptional regulatory protein AlgB |
| **KCPMINPF_16284** | Alkaline phosphatase synthesis transcriptional regulatory protein SphR |
| **KCPMINPF_162912** | Transcriptional regulator MraZ |
| **KCPMINPF_163010** | Regulatory protein AtoC |
| **KCPMINPF_163053** | Transcriptional regulatory protein WalR |
| **KCPMINPF_163085** | Transcriptional regulatory protein ros |
| **KCPMINPF_163146** | Photosynthetic apparatus regulatory protein RegA |
| **KCPMINPF_163177** | RNA polymerase-binding transcription factor DksA |
| **KCPMINPF_163211** | Mercuric resistance operon regulatory protein |
| **KCPMINPF_163346** | Transcriptional regulatory protein KdpE |
| **KCPMINPF_163466** | Transcriptional regulatory protein YpdB |
| **KCPMINPF_163725** | Transcriptional regulatory protein LnrK |
| **KCPMINPF_164123** | Nitrogen regulatory protein P-II 1 |
| **KCPMINPF_164126** | Transcriptional regulatory protein OmpR |
| **KCPMINPF_164139** | Transcriptional regulatory protein ZraR |
| **KCPMINPF_164159** | Copper-sensing transcriptional repressor CsoR |
| **KCPMINPF_164306** | cAMP-activated global transcriptional regulator CRP |
| **KCPMINPF_164333** | DNA-binding transcriptional activator DecR |
| **KCPMINPF_164337** | Transcriptional regulatory protein CusR |
| **KCPMINPF_164377** | KDP operon transcriptional regulatory protein KdpE |
| **KCPMINPF_164400** | KDP operon transcriptional regulatory protein KdpE |
| **KCPMINPF_164618** | Transcriptional regulatory protein CusR |
| **KCPMINPF_164744** | DNA-binding transcriptional activator DevR/DosR |
| **KCPMINPF_164841** | Transcriptional regulator PerR |
| **KCPMINPF_165520** | LexA repressor |
| **KCPMINPF_165592** | LexA repressor |
| **KCPMINPF_165628** | Regulatory protein AtoC |
| **KCPMINPF_165872** | Transcriptional regulatory protein OmpR |
| **KCPMINPF_165904** | Nitrogen regulatory protein |
| **KCPMINPF_166015** | Flagellar transcriptional regulator FtcR |
| **KCPMINPF_16670** | flagellum biosynthesis repressor protein FlbT |
| **KCPMINPF_16678** | RNA polymerase-binding transcription factor DksA |
| **KCPMINPF_16768** | Transcriptional repressor PaaX |
| **KCPMINPF_16780** | Regulatory protein PchR |
| **KCPMINPF_17001** | Regulatory protein RecX |
| **KCPMINPF_17046** | Bifunctional ligase/repressor BirA |
| **KCPMINPF_17326** | Alkaline phosphatase synthesis transcriptional regulatory protein PhoP |

| | |
|---|---|
| **KCPMINPF_17429** | Transcriptional regulatory protein LiaR |
| **KCPMINPF_17503** | Bifunctional ligase/repressor BirA |
| **KCPMINPF_17644** | Transcriptional regulatory protein ZraR |
| **KCPMINPF_17658** | Regulatory protein AsnC |
| **KCPMINPF_17663** | Regulatory protein RecX |
| **KCPMINPF_17759** | Phosphate regulon transcriptional regulatory protein PhoB |
| **KCPMINPF_17847** | Nitrogen regulatory protein P-II |
| **KCPMINPF_17928** | Transcriptional regulatory protein BtsR |
| **KCPMINPF_18004** | Arsenic resistance transcriptional regulator ArsR2 |
| **KCPMINPF_18212** | RNA polymerase-binding transcription factor DksA |
| **KCPMINPF_18326** | Transcriptional regulatory protein QseB |
| **KCPMINPF_18336** | Transcriptional regulatory protein CpxR |
| **KCPMINPF_18438** | Arginine repressor |
| **KCPMINPF_18446** | Transcriptional regulator MntR |
| **KCPMINPF_18520** | LexA repressor |
| **KCPMINPF_18556** | Bifunctional ligase/repressor BirA |
| **KCPMINPF_18617** | Transcriptional regulatory protein DegU |
| **KCPMINPF_18623** | RNA polymerase-binding transcription factor DksA |
| **KCPMINPF_18711** | Transcriptional repressor NrdR |
| **KCPMINPF_18725** | Leucine-responsive regulatory protein |
| **KCPMINPF_18970** | Transcriptional regulatory protein ZraR |
| **KCPMINPF_19003** | Transcriptional regulator MraZ |
| **KCPMINPF_19005** | Oxygen regulatory protein NreC |
| **KCPMINPF_19069** | Transcriptional regulatory protein ZraR |
| **KCPMINPF_19113** | LexA repressor |
| **KCPMINPF_19149** | Regulatory protein AtoC |
| **KCPMINPF_19167** | Transcriptional regulatory protein ZraR |
| **KCPMINPF_19183** | Transcriptional regulatory protein ZraR |
| **KCPMINPF_19254** | Nitrogen regulatory protein P-II 2 |
| **KCPMINPF_19295** | Transcriptional regulatory protein QseB |
| **KCPMINPF_19364** | Transcriptional regulatory protein KdpE |
| **KCPMINPF_19376** | Nitrogen regulatory protein P-II 2 |
| **KCPMINPF_19421** | KDP operon transcriptional regulatory protein KdpE |
| **KCPMINPF_19442** | Transcriptional regulatory protein tctD |
| **KCPMINPF_19588** | Fumarate and nitrate reduction regulatory protein |
| **KCPMINPF_19592** | Transcriptional regulatory protein BasR |
| **KCPMINPF_19600** | DnaA regulatory inactivator Hda |
| **KCPMINPF_19699** | Regulatory protein AtoC |
| **KCPMINPF_19910** | Heat-inducible transcription repressor HrcA |
| **KCPMINPF_19947** | Transcriptional regulator WhiB |
| **KCPMINPF_19962** | Transcriptional regulator WhiB |
| **KCPMINPF_20128** | LexA repressor |
| **KCPMINPF_20209** | Transcriptional repressor NrdR |

| | |
|---|---|
| **KCPMINPF_20269** | Formate hydrogenlyase transcriptional activator FhlA |
| **KCPMINPF_20497** | Octopine catabolism/uptake operon regulatory protein OccR |
| **KCPMINPF_20512** | Mannosyl-D-glycerate transport/metabolism system repressor MngR |
| **KCPMINPF_20568** | N-acetylglucosamine repressor |
| **KCPMINPF_20569** | Phosphate regulon transcriptional regulatory protein PhoB |
| **KCPMINPF_20582** | C4-dicarboxylate transport transcriptional regulatory protein DctD |
| **KCPMINPF_20605** | DNA-binding transcriptional regulator BolA |
| **KCPMINPF_20742** | Regulatory protein AtoC |
| **KCPMINPF_21072** | Transcriptional repressor NrdR |
| **KCPMINPF_21089** | Transcriptional regulator MraZ |
| **KCPMINPF_21156** | Transcriptional regulatory protein WalR |
| **KCPMINPF_21279** | Transcriptional repressor SmtB |
| **KCPMINPF_21282** | Transcriptional regulatory protein BaeR |
| **KCPMINPF_21348** | Transcriptional regulatory protein DegU |
| **KCPMINPF_21434** | Transcriptional regulatory protein LiaR |
| **KCPMINPF_21560** | Regulatory protein AtoC |
| **KCPMINPF_21568** | Transcriptional regulatory protein AfsQ1 |
| **KCPMINPF_21572** | Oxygen regulatory protein NreC |
| **KCPMINPF_21636** | RNA polymerase-binding transcription factor DksA |
| **KCPMINPF_21815** | Regulatory protein AtoC |
| **KCPMINPF_21817** | RNA polymerase-binding transcription factor DksA |
| **KCPMINPF_21837** | Transcriptional regulatory protein CreB |
| **KCPMINPF_21934** | Aspartate carbamoyltransferase regulatory chain |
| **KCPMINPF_21974** | Regulatory protein AtoC |
| **KCPMINPF_22046** | Regulatory protein AtoC |
| **KCPMINPF_22131** | Glucitol operon repressor |
| **KCPMINPF_22202** | Ribose operon repressor |
| **KCPMINPF_22238** | Glucitol operon repressor |
| **KCPMINPF_22307** | LexA repressor |
| **KCPMINPF_22321** | Transcriptional regulator KdgR |
| **KCPMINPF_22353** | Transcriptional regulatory protein LiaR |
| **KCPMINPF_22573** | Transcriptional regulatory protein DegU |
| **KCPMINPF_22693** | Transcriptional regulatory protein WalR |
| **KCPMINPF_22729** | Transcriptional regulatory protein ZraR |
| **KCPMINPF_22733** | Regulatory protein AtoC |
| **KCPMINPF_22798** | Alkaline phosphatase synthesis transcriptional regulatory protein PhoP |
| **KCPMINPF_22886** | LexA repressor |
| **KCPMINPF_23065** | Alkaline phosphatase synthesis transcriptional regulatory protein PhoP |
| **KCPMINPF_23121** | Transcriptional regulatory protein LnrK |
| **KCPMINPF_23151** | p-hydroxybenzoate hydroxylase transcriptional activator |
| **KCPMINPF_23173** | Fumarate and nitrate reduction regulatory protein |
| **KCPMINPF_23272** | Transcriptional regulatory protein SrrA |
| **KCPMINPF_23635** | Zinc-specific metallo-regulatory protein |

| | |
|---|---|
| **KCPMINPF_23752** | Oxygen regulatory protein NreC |
| **KCPMINPF_23774** | Transcriptional regulatory protein YpdB |
| **KCPMINPF_23873** | Transcriptional regulatory protein ZraR |
| **KCPMINPF_23917** | Transcriptional regulatory protein BtsR |
| **KCPMINPF_24170** | Regulatory protein AtoC |
| **KCPMINPF_24222** | LexA repressor |
| **KCPMINPF_24243** | Alkaline phosphatase synthesis transcriptional regulatory protein PhoP |
| **KCPMINPF_24418** | DNA-binding transcriptional activator DecR |
| **KCPMINPF_24657** | Regulatory protein AtoC |
| **KCPMINPF_24709** | Transcriptional regulatory protein ZraR |
| **KCPMINPF_24827** | C4-dicarboxylate transport transcriptional regulatory protein DctD |
| **KCPMINPF_24907** | Regulatory protein AsnC |
| **KCPMINPF_25063** | Regulatory protein AtoC |
| **KCPMINPF_25086** | Alkaline phosphatase synthesis transcriptional regulatory protein PhoP |
| **KCPMINPF_25104** | Transcriptional regulatory protein DegU |
| **KCPMINPF_25133** | Redox-sensing transcriptional repressor Rex |
| **KCPMINPF_25381** | Transcriptional regulatory protein SrrA |
| **KCPMINPF_25476** | Phosphate regulon transcriptional regulatory protein PhoB |
| **KCPMINPF_25532** | Bifunctional ligase/repressor BirA |
| **KCPMINPF_25554** | Regulatory protein AtoC |
| **KCPMINPF_25555** | Regulatory protein AtoC |
| **KCPMINPF_25574** | Regulatory protein AtoC |
| **KCPMINPF_25628** | Heat-inducible transcription repressor HrcA |
| **KCPMINPF_25801** | Transcriptional regulatory protein OmpR |
| **KCPMINPF_25825** | Leucine-responsive regulatory protein |
| **KCPMINPF_25836** | Transcriptional regulator SlyA |
| **KCPMINPF_25992** | Redox-sensing transcriptional repressor Rex |
| **KCPMINPF_26221** | Arginine repressor |
| **KCPMINPF_26543** | Transcriptional activator protein CopR |
| **KCPMINPF_26745** | Diphtheria toxin repressor |
| **KCPMINPF_26773** | Transcriptional regulatory protein BtsR |
| **KCPMINPF_26895** | Transcriptional regulator WhiD |
| **KCPMINPF_26991** | Iron-dependent repressor IdeR |
| **KCPMINPF_26996** | Nitrogen regulatory protein P-II |
| **KCPMINPF_27010** | Transcriptional regulator WhiB |
| **KCPMINPF_27169** | Transcriptional regulatory protein OmpR |
| **KCPMINPF_27203** | Fumarate and nitrate reduction regulatory protein |
| **KCPMINPF_27447** | Alkaline phosphatase synthesis transcriptional regulatory protein PhoP |
| **KCPMINPF_27448** | Alkaline phosphatase synthesis transcriptional regulatory protein PhoP |
| **KCPMINPF_27480** | Alkaline phosphatase synthesis transcriptional regulatory protein PhoP |
| **KCPMINPF_27492** | Oxygen regulatory protein NreC |
| **KCPMINPF_27743** | Transcriptional regulatory protein PhoP |
| **KCPMINPF_27751** | Transcriptional regulatory protein CusR |

| | |
|---|---|
| **KCPMINPF_27907** | Bifunctional ligase/repressor BirA |
| **KCPMINPF_28236** | Nitrogen regulatory protein P-II |
| **KCPMINPF_28558** | Mannosyl-D-glycerate transport/metabolism system repressor MngR |
| **KCPMINPF_28624** | Transcriptional regulatory protein FixJ |
| **KCPMINPF_28672** | Hydrogen peroxide-inducible genes activator |
| **KCPMINPF_28714** | LexA repressor |
| **KCPMINPF_28832** | Transcriptional repressor IclR |
| **KCPMINPF_28851** | Transcriptional regulatory protein FixJ |
| **KCPMINPF_28863** | Transcriptional regulatory protein BtsR |
| **KCPMINPF_28874** | Nitrogen regulatory protein P-II |
| **KCPMINPF_29017** | Transcriptional regulatory protein ZraR |
| **KCPMINPF_29166** | Transcriptional regulatory protein LiaR |
| **KCPMINPF_29212** | Methanol dehydrogenase activator |
| **KCPMINPF_29330** | Penicillinase repressor |
| **KCPMINPF_29433** | Alkaline phosphatase synthesis transcriptional regulatory protein PhoP |
| **KCPMINPF_29688** | N-acetylglucosamine repressor |
| **KCPMINPF_29694** | N-acetylglucosamine repressor |
| **KCPMINPF_29784** | Hydrogen peroxide-inducible genes activator |
| **KCPMINPF_29790** | Glycerol-3-phosphate regulon repressor |
| **KCPMINPF_29836** | Pyruvate dehydrogenase complex repressor |
| **KCPMINPF_29840** | ATP phosphoribosyltransferase regulatory subunit |
| **KCPMINPF_29967** | Hydrogen peroxide-inducible genes activator |
| **KCPMINPF_30105** | LexA repressor |
| **KCPMINPF_30164** | Transcriptional regulatory protein KdpE |
| **KCPMINPF_30363** | Transcriptional regulator KdgR |
| **KCPMINPF_30384** | Transcriptional regulatory protein OmpR |
| **KCPMINPF_30427** | Leucine-responsive regulatory protein |
| **KCPMINPF_30519** | Transcriptional regulatory protein QseB |
| **KCPMINPF_30602** | Nitrogen regulatory protein P-II |
| **KCPMINPF_30631** | Regulatory protein AtoC |
| **KCPMINPF_30695** | Alkaline phosphatase synthesis transcriptional regulatory protein PhoP |
| **KCPMINPF_30838** | Transcriptional regulatory protein QseB |
| **KCPMINPF_30893** | Transcriptional regulator AcuR |
| **KCPMINPF_30958** | Transcriptional regulatory protein KdpE |
| **KCPMINPF_31039** | Transcriptional regulatory protein DegU |
| **KCPMINPF_31053** | Transcriptional regulator MraZ |
| **KCPMINPF_31108** | Bifunctional ligase/repressor BirA |
| **KCPMINPF_31246** | Transcriptional repressor NrdR |
| **KCPMINPF_31505** | Transcriptional regulatory protein DegU |
| **KCPMINPF_31626** | Transcriptional activatory protein BadR |
| **KCPMINPF_31686** | DNA-binding transcriptional activator DecR |
| **KCPMINPF_31731** | Transcriptional regulatory protein BaeR |
| **KCPMINPF_31832** | Transcriptional regulatory protein WalR |

| | |
|---|---|
| **KCPMINPF_31844** | Alkaline phosphatase synthesis transcriptional regulatory protein PhoP |
| **KCPMINPF_31922** | Transcriptional regulatory protein DegU |
| **KCPMINPF_31963** | Regulatory protein AtoC |
| **KCPMINPF_32046** | N-acetylglucosamine repressor |
| **KCPMINPF_32287** | Transcriptional regulatory protein CusR |
| **KCPMINPF_32291** | Alkaline phosphatase synthesis transcriptional regulatory protein PhoP |
| **KCPMINPF_32311** | Protein-arginine kinase activator protein |
| **KCPMINPF_32541** | Alkaline phosphatase synthesis transcriptional regulatory protein PhoP |
| **KCPMINPF_32690** | Transcriptional regulatory protein WalR |
| **KCPMINPF_32699** | Copper-sensing transcriptional repressor CsoR |
| **KCPMINPF_32758** | Transcriptional regulator WhiB |
| **KCPMINPF_32771** | Transcriptional regulatory protein DegU |
| **KCPMINPF_32777** | Hydrogen peroxide-inducible genes activator |
| **KCPMINPF_32886** | Transcriptional regulator SlyA |
| **KCPMINPF_32890** | Transcriptional regulatory protein BtsR |
| **KCPMINPF_32950** | Transcriptional regulatory protein DegU |
| **KCPMINPF_33034** | Heat-inducible transcription repressor HrcA |
| **KCPMINPF_33050** | Transcriptional repressor NrdR |
| **KCPMINPF_33057** | Regulatory protein AtoC |
| **KCPMINPF_33086** | Arginine repressor |
| **KCPMINPF_33202** | Regulatory protein AsnC |
| **KCPMINPF_33254** | Iron-dependent repressor IdeR |
| **KCPMINPF_33345** | Transcriptional regulator MraZ |
| **KCPMINPF_33556** | Fumarate and nitrate reduction regulatory protein |
| **KCPMINPF_33636** | Alkaline phosphatase synthesis transcriptional regulatory protein SphR |
| **KCPMINPF_33824** | Formate hydrogenlyase transcriptional activator FhlA |
| **KCPMINPF_33844** | Regulatory protein AtoC |
| **KCPMINPF_33847** | Regulatory protein AtoC |
| **KCPMINPF_33886** | Lactose operon repressor |
| **KCPMINPF_33899** | Transcriptional regulatory protein LiaR |
| **KCPMINPF_34023** | Heat-inducible transcription repressor HrcA |
| **KCPMINPF_34042** | Phosphoenolpyruvate synthase regulatory protein |
| **KCPMINPF_34054** | Glycine cleavage system transcriptional repressor |
| **KCPMINPF_34139** | DnaA regulatory inactivator Hda |
| **KCPMINPF_34414** | Bifunctional ligase/repressor BirA |
| **KCPMINPF_34513** | Murein hydrolase activator EnvC |
| **KCPMINPF_34534** | Regulatory protein AtoC |
| **KCPMINPF_34682** | LexA repressor |
| **KCPMINPF_34757** | LexA repressor |
| **KCPMINPF_34997** | Transcriptional activator protein Anr |
| **KCPMINPF_35009** | Leucine-responsive regulatory protein |
| **KCPMINPF_35030** | Transcriptional activator NphR |
| **KCPMINPF_35059** | Glycine cleavage system transcriptional activator |

| | |
|---|---|
| **KCPMINPF_35078** | Transcriptional regulatory protein QseB |
| **KCPMINPF_35238** | Methanol dehydrogenase activator |
| **KCPMINPF_35554** | Transcriptional regulatory protein LiaR |
| **KCPMINPF_35665** | Nitrogen regulatory protein P-II |
| **KCPMINPF_35763** | Transcriptional regulatory protein LiaR |
| **KCPMINPF_35852** | LexA repressor |
| **KCPMINPF_35901** | Heat-inducible transcription repressor HrcA |
| **KCPMINPF_35935** | Regulatory protein RecX |
| **KCPMINPF_35951** | Copper-sensing transcriptional repressor CsoR |
| **KCPMINPF_36159** | Leucine-responsive regulatory protein |
| **KCPMINPF_36201** | Biofilm growth-associated repressor |
| **KCPMINPF_36225** | Pca regulon regulatory protein |
| **KCPMINPF_36226** | Transcriptional regulator KdgR |
| **KCPMINPF_36374** | Transcriptional regulatory protein WalR |
| **KCPMINPF_36406** | LexA repressor |
| **KCPMINPF_36628** | Bifunctional transcriptional activator/DNA repair enzyme Ada |
| **KCPMINPF_36849** | Alkaline phosphatase synthesis transcriptional regulatory protein SphR |
| **KCPMINPF_36942** | DNA-binding transcriptional activator EvgA |
| **KCPMINPF_36979** | RNA polymerase-binding transcription factor DksA |
| **KCPMINPF_36984** | Hydrogen peroxide-inducible genes activator |
| **KCPMINPF_37021** | Regulatory protein AtoC |
| **KCPMINPF_37060** | PCP degradation transcriptional activation protein |
| **KCPMINPF_37168** | Alkaline phosphatase synthesis transcriptional regulatory protein PhoP |
| **KCPMINPF_37191** | Transcriptional regulatory protein OmpR |
| **KCPMINPF_37453** | Transcriptional regulatory protein TcrA |
| **KCPMINPF_37526** | Methanol dehydrogenase activator |
| **KCPMINPF_37612** | Regulatory protein AtoC |
| **KCPMINPF_37742** | Aspartate carbamoyltransferase regulatory chain |
| **KCPMINPF_37823** | Transcriptional regulatory protein LiaR |
| **KCPMINPF_38093** | Heat-inducible transcription repressor HrcA |
| **KCPMINPF_38173** | Transcriptional regulator MraZ |
| **KCPMINPF_38255** | Methanol dehydrogenase activator |
| **KCPMINPF_38287** | DNA-binding transcriptional regulator NtrC |
| **KCPMINPF_38520** | Transcriptional repressor IclR |
| **KCPMINPF_38522** | Transcriptional regulator KdgR |
| **KCPMINPF_38652** | Murein hydrolase activator EnvC |
| **KCPMINPF_38671** | KDP operon transcriptional regulatory protein KdpE |
| **KCPMINPF_38739** | Anaerobic regulatory protein |
| **KCPMINPF_38794** | Murein hydrolase activator EnvC |
| **KCPMINPF_38910** | Heat-inducible transcription repressor HrcA |
| **KCPMINPF_39079** | Heat-inducible transcription repressor HrcA |
| **KCPMINPF_39098** | Photosynthetic apparatus regulatory protein RegA |
| **KCPMINPF_39212** | Regulatory protein AtoC |

| | |
|---|---|
| **KCPMINPF_39333** | Regulatory protein AsnC |
| **KCPMINPF_39564** | Transcriptional regulator SlyA |
| **KCPMINPF_39726** | Transcriptional regulatory protein OmpR |
| **KCPMINPF_39828** | Transcriptional regulatory protein DegU |
| **KCPMINPF_39855** | Copper-sensing transcriptional repressor CsoR |
| **KCPMINPF_39872** | Transcriptional regulatory protein DegU |
| **KCPMINPF_39916** | Alkaline phosphatase synthesis transcriptional regulatory protein SphR |
| **KCPMINPF_39932** | Transcriptional repressor SmtB |
| **KCPMINPF_40000** | Regulatory protein RecX |
| **KCPMINPF_40096** | Transcriptional repressor NrdR |
| **KCPMINPF_40104** | Transcriptional regulator MraZ |
| **KCPMINPF_40136** | RNA polymerase-binding transcription factor CarD |
| **KCPMINPF_40203** | Leucine-responsive regulatory protein |
| **KCPMINPF_40415** | Transcriptional regulatory protein WalR |
| **KCPMINPF_40536** | Transcriptional repressor NrdR |
| **KCPMINPF_40563** | Alkaline phosphatase synthesis transcriptional regulatory protein PhoP |
| **KCPMINPF_40646** | Transcriptional regulatory protein DegU |
| **KCPMINPF_40845** | ATP phosphoribosyltransferase regulatory subunit |
| **KCPMINPF_40886** | Transcriptional regulatory protein WalR |
| **KCPMINPF_41011** | LexA repressor |
| **KCPMINPF_41015** | Regulatory protein AtoC |
| **KCPMINPF_41018** | Transcriptional regulatory protein LnrK |
| **KCPMINPF_41066** | Leucine-responsive regulatory protein |
| **KCPMINPF_41126** | LexA repressor |
| **KCPMINPF_41252** | Methanol dehydrogenase activator |
| **KCPMINPF_41338** | Transcriptional regulator LsrR |
| **KCPMINPF_41359** | Transcriptional regulatory protein AfsQ1 |
| **KCPMINPF_41495** | Alkaline phosphatase synthesis transcriptional regulatory protein PhoP |
| **KCPMINPF_41507** | Transcriptional regulator SlyA |
| **KCPMINPF_41567** | Acetoin catabolism regulatory protein |
| **KCPMINPF_41594** | Alkaline phosphatase synthesis transcriptional regulatory protein SphR |
| **KCPMINPF_41654** | DNA-binding transcriptional activator DevR/DosR |
| **KCPMINPF_41655** | Transcriptional regulatory protein WalR |
| **KCPMINPF_41742** | Arginine repressor |
| **KCPMINPF_41798** | Murein hydrolase activator EnvC |
| **KCPMINPF_41982** | cAMP-activated global transcriptional regulator CRP |
| **KCPMINPF_42083** | Nitrogen regulatory protein P-II |
| **KCPMINPF_42111** | Bifunctional ligase/repressor BirA |
| **KCPMINPF_42141** | LexA repressor |
| **KCPMINPF_42340** | Biofilm regulatory protein A |
| **KCPMINPF_42342** | Transcriptional activator protein CopR |
| **KCPMINPF_42352** | Zinc-specific metallo-regulatory protein |
| **KCPMINPF_42412** | Regulatory protein AtoC |

| | |
|---|---|
| **KCPMINPF_42427** | Transcriptional regulator MraZ |
| **KCPMINPF_42508** | Regulatory protein AtoC |
| **KCPMINPF_42729** | Transcriptional regulatory protein KdpE |
| **KCPMINPF_42738** | Transcriptional regulatory protein WalR |
| **KCPMINPF_42849** | Regulatory protein AtoC |
| **KCPMINPF_42932** | Regulatory protein AsnC |
| **KCPMINPF_43013** | Transcriptional regulatory protein ZraR |
| **KCPMINPF_43385** | Transcriptional regulatory protein LiaR |
| **KCPMINPF_43401** | Alkaline phosphatase synthesis transcriptional regulatory protein PhoP |
| **KCPMINPF_43408** | Oxygen regulatory protein NreC |
| **KCPMINPF_43443** | Alkaline phosphatase synthesis transcriptional regulatory protein SphR |
| **KCPMINPF_43513** | Iron-dependent repressor IdeR |
| **KCPMINPF_43599** | Regulatory protein RecX |
| **KCPMINPF_43641** | Iron-dependent repressor IdeR |
| **KCPMINPF_43669** | N-acetylglucosamine repressor |
| **KCPMINPF_43730** | Transcriptional regulatory protein BaeR |
| **KCPMINPF_43747** | Transcriptional regulatory protein WalR |
| **KCPMINPF_43850** | Phosphate regulon transcriptional regulatory protein PhoB |
| **KCPMINPF_43868** | LexA repressor |
| **KCPMINPF_43896** | Transcriptional regulatory protein DegU |
| **KCPMINPF_43949** | Nitrogen regulatory protein |
| **KCPMINPF_43964** | Biofilm growth-associated repressor |
| **KCPMINPF_44024** | Anaerobic regulatory protein |
| **KCPMINPF_44306** | Alkaline phosphatase synthesis transcriptional regulatory protein SphR |
| **KCPMINPF_44355** | Copper-sensing transcriptional repressor CsoR |
| **KCPMINPF_44581** | Oxygen regulatory protein NreC |
| **KCPMINPF_44598** | Transcriptional regulatory protein DegU |
| **KCPMINPF_44665** | Transcriptional regulator SlyA |
| **KCPMINPF_44836** | Transcriptional regulatory protein LiaR |
| **KCPMINPF_44908** | DNA-binding transcriptional activator DevR/DosR |
| **KCPMINPF_44913** | Purine catabolism regulatory protein |
| **KCPMINPF_45017** | Bifunctional ligase/repressor BirA |
| **KCPMINPF_45308** | Copper-sensing transcriptional repressor CsoR |
| **KCPMINPF_45498** | Bifunctional ligase/repressor BirA |
| **KCPMINPF_45575** | Regulatory protein AtoC |
| **KCPMINPF_45946** | Heat-inducible transcription repressor HrcA |
| **KCPMINPF_45993** | Phosphate regulon transcriptional regulatory protein PhoB |
| **KCPMINPF_46066** | Leucine-responsive regulatory protein |
| **KCPMINPF_46070** | RNA polymerase-binding transcription factor DksA |
| **KCPMINPF_46141** | Transcriptional regulatory protein tctD |
| **KCPMINPF_46376** | Transcriptional regulatory protein FixJ |
| **KCPMINPF_46517** | Anaerobic regulatory protein |
| **KCPMINPF_46591** | Bifunctional ligase/repressor BirA |

| | |
|---|---|
| **KCPMINPF_46631** | Transcriptional regulator PerR |
| **KCPMINPF_46953** | Lactose operon repressor |
| **KCPMINPF_46974** | Glycerol-3-phosphate regulon repressor |
| **KCPMINPF_47012** | Regulatory protein AtoC |
| **KCPMINPF_47035** | Regulatory protein AtoC |
| **KCPMINPF_47219** | Hydrogen peroxide-inducible genes activator |
| **KCPMINPF_47223** | Glycine cleavage system transcriptional activator |
| **KCPMINPF_47241** | Transcriptional activator HlyU |
| **KCPMINPF_47292** | Transcriptional regulatory protein WalR |
| **KCPMINPF_47295** | Transcriptional regulatory protein QseB |
| **KCPMINPF_47307** | Transcriptional regulator SlyA |
| **KCPMINPF_47311** | Transcriptional regulatory protein DegU |
| **KCPMINPF_47465** | Transcriptional regulator MraZ |
| **KCPMINPF_47492** | Alkaline phosphatase synthesis transcriptional regulatory protein PhoP |
| **KCPMINPF_47598** | Copper-sensing transcriptional repressor CsoR |
| **KCPMINPF_47802** | Phosphate regulon transcriptional regulatory protein PhoB |
| **KCPMINPF_47803** | DNA-binding transcriptional activator DevR/DosR |
| **KCPMINPF_47865** | Transcriptional regulatory protein LnrK |
| **KCPMINPF_47954** | cAMP-activated global transcriptional regulator CRP |
| **KCPMINPF_47958** | Nitrogen regulatory protein P-II |
| **KCPMINPF_47992** | Transcriptional activator protein Anr |
| **KCPMINPF_48156** | Transcriptional regulatory protein DegU |
| **KCPMINPF_48199** | Regulatory protein MsrR |
| **KCPMINPF_48252** | Glycerol-3-phosphate regulon repressor |
| **KCPMINPF_48340** | KDP operon transcriptional regulatory protein KdpE |
| **KCPMINPF_48377** | Nitrogen regulatory protein P-II |
| **KCPMINPF_48423** | N-acetylglucosamine repressor |
| **KCPMINPF_48452** | Transcriptional regulator MraZ |
| **KCPMINPF_48548** | Transcriptional regulatory protein WalR |
| **KCPMINPF_48570** | Alkaline phosphatase synthesis transcriptional regulatory protein PhoP |
| **KCPMINPF_48651** | Alkaline phosphatase synthesis transcriptional regulatory protein PhoP |
| **KCPMINPF_48683** | Regulatory protein AtoC |
| **KCPMINPF_48704** | Transcriptional regulatory protein ZraR |
| **KCPMINPF_48716** | Transcriptional regulatory protein PmpR |
| **KCPMINPF_48778** | LexA repressor |
| **KCPMINPF_48887** | Bifunctional ligase/repressor BirA |
| **KCPMINPF_49066** | Regulatory protein AtoC |
| **KCPMINPF_49127** | Transcriptional repressor NrdR |
| **KCPMINPF_49139** | Regulatory protein AtoC |
| **KCPMINPF_49164** | Regulatory protein RecX |
| **KCPMINPF_49460** | Regulatory protein AtoC |
| **KCPMINPF_49467** | Alkaline phosphatase synthesis transcriptional regulatory protein PhoP |
| **KCPMINPF_49607** | Transcriptional regulator LsrR |

| | |
|---|---|
| **KCPMINPF_49661** | RNA polymerase-binding transcription factor DksA |
| **KCPMINPF_49689** | DNA-binding transcriptional regulator NtrC |
| **KCPMINPF_49694** | Alkaline phosphatase synthesis transcriptional regulatory protein PhoP |
| **KCPMINPF_49706** | Redox-sensing transcriptional repressor Rex |
| **KCPMINPF_49770** | Transcriptional regulatory protein KdpE |
| **KCPMINPF_49792** | Transcriptional regulatory protein QseB |
| **KCPMINPF_49826** | DNA-binding transcriptional activator DecR |
| **KCPMINPF_49858** | Transcriptional activator HlyU |
| **KCPMINPF_50095** | Glucitol operon repressor |
| **KCPMINPF_50101** | Transcriptional regulator ManR |
| **KCPMINPF_50202** | Transcriptional repressor NrdR |
| **KCPMINPF_50337** | RNA polymerase-binding transcription factor DksA |
| **KCPMINPF_50509** | Transcriptional repressor SmtB |
| **KCPMINPF_50524** | Transcriptional regulatory protein LiaR |
| **KCPMINPF_50535** | Regulatory protein AtoC |
| **KCPMINPF_50537** | Oxygen regulatory protein NreC |
| **KCPMINPF_50587** | Glucitol operon repressor |
| **KCPMINPF_50675** | Alkaline phosphatase synthesis transcriptional regulatory protein PhoP |
| **KCPMINPF_50785** | RNA polymerase-binding transcription factor DksA |
| **KCPMINPF_50815** | Transcriptional regulatory protein BtsR |
| **KCPMINPF_50836** | Nif-specific regulatory protein |
| **KCPMINPF_51012** | Transcriptional regulatory protein WalR |
| **KCPMINPF_51215** | Transcriptional regulator SlyA |
| **KCPMINPF_51230** | RNA polymerase-binding transcription factor DksA |
| **KCPMINPF_51249** | Octopine catabolism/uptake operon regulatory protein OccR |
| **KCPMINPF_51259** | Transcriptional regulator HilA |
| **KCPMINPF_51284** | (R)-phenyllactate dehydratase activator |
| **KCPMINPF_51285** | (R)-phenyllactate dehydratase activator |
| **KCPMINPF_51500** | Transcriptional regulatory protein LiaR |
| **KCPMINPF_51631** | Iron-dependent repressor IdeR |
| **KCPMINPF_51926** | Alkaline phosphatase synthesis transcriptional regulatory protein PhoP |
| **KCPMINPF_52055** | Transcriptional regulatory protein CusR |
| **KCPMINPF_52185** | Transcriptional regulatory protein LiaR |
| **KCPMINPF_52205** | Bifunctional transcriptional activator/DNA repair enzyme Ada |
| **KCPMINPF_52240** | Redox-sensing transcriptional repressor Rex |
| **KCPMINPF_52245** | Redox-sensing transcriptional repressor Rex |
| **KCPMINPF_52268** | Bifunctional ligase/repressor BirA |
| **KCPMINPF_52299** | Oxygen regulatory protein NreC |
| **KCPMINPF_52408** | Transcriptional regulator MraZ |
| **KCPMINPF_52683** | Acetoin catabolism regulatory protein |
| **KCPMINPF_52734** | RNA polymerase-binding transcription factor DksA |
| **KCPMINPF_52764** | Transcriptional regulator MraZ |
| **KCPMINPF_52838** | Regulatory protein AtoC |

| | |
|---|---|
| **KCPMINPF_52848** | Heat-inducible transcription repressor HrcA |
| **KCPMINPF_52898** | Regulatory protein AtoC |
| **KCPMINPF_52908** | Heat-inducible transcription repressor HrcA |
| **KCPMINPF_53053** | PCP degradation transcriptional activation protein |
| **KCPMINPF_53068** | Transcriptional regulatory protein WalR |
| **KCPMINPF_53076** | Nitrogen regulatory protein P-II 2 |
| **KCPMINPF_53078** | Nitrogen regulatory protein P-II 2 |
| **KCPMINPF_53172** | Transcriptional regulatory protein QseF |
| **KCPMINPF_53190** | Penicillin-binding protein activator LpoB |
| **KCPMINPF_53256** | Transcriptional regulator WhiB2 |
| **KCPMINPF_53263** | Transcriptional regulator WhiB |
| **KCPMINPF_53403** | Regulatory protein AtoC |
| **KCPMINPF_53461** | Transcriptional regulatory protein DegU |
| **KCPMINPF_53726** | Photosynthetic apparatus regulatory protein RegA |
| **KCPMINPF_53751** | Transcriptional regulatory protein DegU |
| **KCPMINPF_54076** | RNA polymerase-binding transcription factor DksA |
| **KCPMINPF_54103** | Phosphoenolpyruvate synthase regulatory protein |
| **KCPMINPF_54189** | 2-hydroxyisocaproyl-CoA dehydratase activator |
| **KCPMINPF_54190** | 2-hydroxyisocaproyl-CoA dehydratase activator |
| **KCPMINPF_54195** | Transcriptional repressor PaaX |
| **KCPMINPF_54450** | Transcriptional regulatory protein BtsR |
| **KCPMINPF_54522** | DNA-binding transcriptional regulator NtrC |
| **KCPMINPF_54556** | Regulatory protein RecX |
| **KCPMINPF_54579** | Transcriptional regulator MraZ |
| **KCPMINPF_54887** | Transcriptional regulatory protein LiaR |
| **KCPMINPF_54900** | ATP phosphoribosyltransferase regulatory subunit |
| **KCPMINPF_54941** | Transcriptional regulatory protein LiaR |
| **KCPMINPF_55043** | Transcriptional regulatory protein WalR |
| **KCPMINPF_55096** | Alkaline phosphatase synthesis transcriptional regulatory protein SphR |
| **KCPMINPF_55191** | Glycerol operon regulatory protein |
| **KCPMINPF_55311** | Transcriptional regulatory protein LiaR |
| **KCPMINPF_55327** | Transcriptional regulatory protein DegU |
| **KCPMINPF_55361** | Iron-dependent repressor IdeR |
| **KCPMINPF_55375** | Alkaline phosphatase synthesis transcriptional regulatory protein PhoP |
| **KCPMINPF_55378** | Transcriptional repressor SmtB |
| **KCPMINPF_55400** | Nitrogen regulatory protein P-II |
| **KCPMINPF_55410** | Bifunctional ligase/repressor BirA |
| **KCPMINPF_55441** | Glycine cleavage system transcriptional activator |
| **KCPMINPF_55448** | DNA-binding transcriptional regulator NtrC |
| **KCPMINPF_55450** | DNA-binding transcriptional regulator NtrC |
| **KCPMINPF_55527** | Leucine-responsive regulatory protein |
| **KCPMINPF_55596** | Photosynthetic apparatus regulatory protein RegA |
| **KCPMINPF_55630** | Pca regulon regulatory protein |

| | |
|---|---|
| **KCPMINPF_55708** | Leucine-responsive regulatory protein |
| **KCPMINPF_55812** | Transcriptional regulator PerR |
| **KCPMINPF_56062** | Heat-inducible transcription repressor HrcA |
| **KCPMINPF_56146** | Alkaline phosphatase synthesis transcriptional regulatory protein PhoP |
| **KCPMINPF_56201** | RNA polymerase-binding transcription factor DksA |
| **KCPMINPF_56372** | Hca operon transcriptional activator HcaR |
| **KCPMINPF_56582** | Transcriptional regulatory protein AfsQ1 |
| **KCPMINPF_56685** | Hydrogen peroxide-inducible genes activator |
| **KCPMINPF_56788** | Transcriptional regulatory protein KdpE |
| **KCPMINPF_56891** | Copper-sensing transcriptional repressor CsoR |
| **KCPMINPF_56910** | RNA polymerase-binding transcription factor DksA |
| **KCPMINPF_56913** | Phosphate regulon transcriptional regulatory protein PhoB |
| **KCPMINPF_57016** | Transcriptional regulatory protein BaeR |
| **KCPMINPF_57058** | Alkaline phosphatase synthesis transcriptional regulatory protein PhoP |
| **KCPMINPF_57063** | Alkaline phosphatase synthesis transcriptional regulatory protein PhoP |
| **KCPMINPF_57095** | Regulatory protein RecX |
| **KCPMINPF_57157** | Transcriptional regulatory protein LnrK |
| **KCPMINPF_57165** | Iron-dependent repressor IdeR |
| **KCPMINPF_57247** | Penicillin-binding protein activator LpoA |
| **KCPMINPF_57252** | Heat-inducible transcription repressor HrcA |
| **KCPMINPF_57302** | ATP phosphoribosyltransferase regulatory subunit |
| **KCPMINPF_57375** | Regulatory protein AtoC |
| **KCPMINPF_57454** | Transcriptional regulator MraZ |
| **KCPMINPF_57540** | Nitrogen regulatory protein P-II 2 |
| **KCPMINPF_57643** | Transcriptional regulatory protein TcrA |
| **KCPMINPF_57648** | Transcriptional regulatory protein QseB |
| **KCPMINPF_57688** | Transcriptional regulatory protein SrrA |
| **KCPMINPF_57725** | Oxygen regulatory protein NreC |
| **KCPMINPF_57910** | Transcriptional repressor NrdR |
| **KCPMINPF_58026** | Transcriptional regulatory protein QseB |
| **KCPMINPF_58180** | N-acetylglucosamine repressor |
| **KCPMINPF_58313** | Transcriptional regulatory protein LiaR |
| **KCPMINPF_58380** | Alkaline phosphatase synthesis transcriptional regulatory protein PhoP |
| **KCPMINPF_58469** | Regulatory protein RecX |
| **KCPMINPF_58497** | DNA-binding transcriptional activator DecR |
| **KCPMINPF_58509** | Transcriptional regulator SlyA |
| **KCPMINPF_58603** | Phosphate regulon transcriptional regulatory protein PhoB |
| **KCPMINPF_58692** | Alkaline phosphatase synthesis transcriptional regulatory protein PhoP |
| **KCPMINPF_58828** | Transcriptional regulatory protein BtsR |
| **KCPMINPF_58868** | Denitrification regulatory protein NirQ |
| **KCPMINPF_58941** | Transcriptional regulatory protein AfsQ1 |
| **KCPMINPF_59133** | Transcriptional regulatory protein DegU |
| **KCPMINPF_59283** | Regulatory protein RecX |

| | |
|---|---|
| **KCPMINPF_59286** | Transcriptional regulator MraZ |
| **KCPMINPF_59303** | Transcriptional repressor NrdR |
| **KCPMINPF_59349** | Transcriptional regulatory protein TcrA |
| **KCPMINPF_59519** | Transcriptional regulatory protein DegU |
| **KCPMINPF_59556** | Murein hydrolase activator EnvC |
| **KCPMINPF_59588** | Bifunctional ligase/repressor BirA |
| **KCPMINPF_59678** | N-acetylglucosamine repressor |
| **KCPMINPF_59981** | Iron-dependent repressor IdeR |
| **KCPMINPF_60032** | Transcriptional regulatory protein LnrK |
| **KCPMINPF_60061** | Transcriptional regulatory protein DesR |
| **KCPMINPF_60194** | Transcriptional repressor IclR |
| **KCPMINPF_60312** | RNA polymerase-binding transcription factor CarD |
| **KCPMINPF_60460** | Transcriptional regulator KdgR |
| **KCPMINPF_60468** | DNA-binding transcriptional regulator BolA |
| **KCPMINPF_60737** | Nitrogen regulatory protein |
| **KCPMINPF_60972** | Transcriptional regulatory protein WalR |
| **KCPMINPF_61337** | Heat-inducible transcription repressor HrcA |
| **KCPMINPF_61481** | Alkaline phosphatase synthesis transcriptional regulatory protein PhoP |
| **KCPMINPF_61691** | Transcriptional regulatory protein QseB |
| **KCPMINPF_61821** | Mercuric resistance operon regulatory protein |
| **KCPMINPF_61828** | Transcriptional activator protein CopR |
| **KCPMINPF_61836** | Transcriptional activator protein CopR |
| **KCPMINPF_61846** | Transcriptional regulatory protein LnrK |
| **KCPMINPF_61914** | RNA polymerase-binding transcription factor DksA |
| **KCPMINPF_62036** | Transcriptional regulatory protein WalR |
| **KCPMINPF_62088** | Transcriptional regulatory protein BtsR |
| **KCPMINPF_62127** | Murein hydrolase activator NlpD |
| **KCPMINPF_62229** | Arabinose metabolism transcriptional repressor |
| **KCPMINPF_62279** | Regulatory protein RecX |
| **KCPMINPF_62293** | Transcriptional regulatory protein WalR |
| **KCPMINPF_62370** | Transcriptional regulator PerR |
| **KCPMINPF_62424** | Transcriptional regulatory protein OmpR |
| **KCPMINPF_62471** | Transcriptional regulatory protein CreB |
| **KCPMINPF_62566** | Transcriptional regulatory protein ZraR |
| **KCPMINPF_62682** | Transcriptional regulatory protein PhoP |
| **KCPMINPF_62691** | Transcriptional regulatory protein CusR |
| **KCPMINPF_63293** | LexA repressor |
| **KCPMINPF_63427** | Transcriptional regulatory protein SrrA |
| **KCPMINPF_63558** | CdaA regulatory protein CdaR |
| **KCPMINPF_63626** | Transcriptional regulatory protein KdpE |
| **KCPMINPF_63648** | Regulatory protein AtoC |
| **KCPMINPF_63844** | Anaerobic regulatory protein |
| **KCPMINPF_63902** | Phosphate regulon transcriptional regulatory protein PhoB |

| | |
|---|---|
| **KCPMINPF_63950** | Bifunctional ligase/repressor BirA |
| **KCPMINPF_63968** | Transcriptional regulatory protein WalR |
| **KCPMINPF_64150** | Glycerol-3-phosphate regulon repressor |
| **KCPMINPF_64202** | Transcriptional regulatory protein LiaR |
| **KCPMINPF_64248** | Hca operon transcriptional activator HcaR |
| **KCPMINPF_64466** | KDP operon transcriptional regulatory protein KdpE |
| **KCPMINPF_64472** | Iron-dependent repressor IdeR |
| **KCPMINPF_64654** | Transcriptional regulatory protein ros |
| **KCPMINPF_64714** | Denitrification regulatory protein NirQ |
| **KCPMINPF_64778** | Nitrogen regulatory protein P-II |
| **KCPMINPF_64892** | N-acetylglucosamine repressor |
| **KCPMINPF_64916** | Transcriptional repressor SmtB |
| **KCPMINPF_64960** | Alkaline phosphatase synthesis transcriptional regulatory protein PhoP |
| **KCPMINPF_64999** | PTS-dependent dihydroxyacetone kinase operon regulatory protein |
| **KCPMINPF_65021** | Alkaline phosphatase synthesis transcriptional regulatory protein SphR |
| **KCPMINPF_65104** | Transcriptional regulatory protein DegU |
| **KCPMINPF_65242** | Transcriptional regulatory protein LiaR |
| **KCPMINPF_65263** | Transcriptional regulator MntR |
| **KCPMINPF_65361** | Hydrogen peroxide-inducible genes activator |
| **KCPMINPF_65553** | Transcriptional regulatory protein DegU |
| **KCPMINPF_65557** | Transcriptional regulatory protein QseB |
| **KCPMINPF_65792** | Sensory/regulatory protein RpfC |
| **KCPMINPF_65796** | Transcriptional repressor NrdR |
| **KCPMINPF_65828** | Bifunctional ligase/repressor BirA |
| **KCPMINPF_65869** | Alkaline phosphatase synthesis transcriptional regulatory protein PhoP |
| **KCPMINPF_66129** | LexA repressor |
| **KCPMINPF_66196** | Pyruvate dehydrogenase complex repressor |
| **KCPMINPF_66388** | N-acetylglucosamine repressor |
| **KCPMINPF_66583** | Transcriptional regulatory protein WalR |
| **KCPMINPF_66620** | Bifunctional ligase/repressor BirA |
| **KCPMINPF_66638** | Transcriptional activator NphR |
| **KCPMINPF_66645** | Penicillin-binding protein activator LpoA |
| **KCPMINPF_66675** | LexA repressor |
| **KCPMINPF_66683** | Extracellular matrix regulatory protein A |
| **KCPMINPF_66794** | Transcriptional regulatory protein DesR |
| **KCPMINPF_67028** | Transcriptional regulatory protein KdpE |
| **KCPMINPF_67156** | DNA-binding transcriptional activator DevR/DosR |
| **KCPMINPF_67161** | Transcriptional regulatory protein WalR |
| **KCPMINPF_67253** | Glucitol operon repressor |
| **KCPMINPF_67301** | Transcriptional regulator SdrP |
| **KCPMINPF_67381** | 2-hydroxyisocaproyl-CoA dehydratase activator |
| **KCPMINPF_67382** | (R)-phenyllactate dehydratase activator |
| **KCPMINPF_67417** | Sigma factor AlgU regulatory protein MucB |

| | |
|---|---|
| **KCPMINPF_67536** | Transcriptional regulatory protein FixJ |
| **KCPMINPF_67774** | Transcriptional repressor NrdR |
| **KCPMINPF_67786** | Transcriptional regulatory protein LnrK |
| **KCPMINPF_67881** | Negative regulatory protein YxlE |
| **KCPMINPF_67972** | Oxygen regulatory protein NreC |
| **KCPMINPF_68032** | Alkaline phosphatase synthesis transcriptional regulatory protein PhoP |
| **KCPMINPF_68116** | Transcriptional regulatory protein DegU |
| **KCPMINPF_68273** | Regulatory protein AtoC |
| **KCPMINPF_68768** | Transcriptional regulatory protein WalR |
| **KCPMINPF_68788** | Alkaline phosphatase synthesis transcriptional regulatory protein PhoP |
| **KCPMINPF_68820** | Transcriptional repressor NrdR |
| **KCPMINPF_68942** | Ribose operon repressor |
| **KCPMINPF_69051** | Transcriptional regulatory protein LiaR |
| **KCPMINPF_69117** | Alkaline phosphatase synthesis transcriptional regulatory protein PhoP |
| **KCPMINPF_69210** | Transcriptional regulatory protein OmpR |
| **KCPMINPF_69293** | Regulatory protein RecX |
| **KCPMINPF_69553** | Glucitol operon repressor |
| **KCPMINPF_69647** | Transcriptional regulatory protein LnrK |
| **KCPMINPF_69779** | Alkaline phosphatase synthesis transcriptional regulatory protein PhoP |
| **KCPMINPF_69799** | Transcriptional activator protein CzcR |
| **KCPMINPF_69935** | Glycerol-3-phosphate regulon repressor |
| **KCPMINPF_70160** | Arginine repressor |
| **KCPMINPF_70217** | LexA repressor |
| **KCPMINPF_70240** | Alkaline phosphatase synthesis transcriptional regulatory protein PhoP |
| **KCPMINPF_70489** | RNA polymerase-binding transcription factor DksA |
| **KCPMINPF_70562** | Transcriptional regulatory protein ZraR |
| **KCPMINPF_70625** | Bifunctional ligase/repressor BirA |
| **KCPMINPF_70627** | Heat-inducible transcription repressor HrcA |
| **KCPMINPF_70645** | Leucine-responsive regulatory protein |
| **KCPMINPF_70842** | Transcriptional regulatory protein WalR |
| **KCPMINPF_70916** | Hydrogen peroxide-inducible genes activator |
| **KCPMINPF_71019** | Regulatory protein AtoC |
| **KCPMINPF_71117** | LexA repressor |
| **KCPMINPF_71143** | Transcriptional regulatory protein KdpE |
| **KCPMINPF_71232** | Nitrogen regulatory protein P-II |
| **KCPMINPF_71332** | (R)-phenyllactate dehydratase activator |
| **KCPMINPF_71333** | (R)-phenyllactate dehydratase activator |
| **KCPMINPF_71469** | Leucine-responsive regulatory protein |
| **KCPMINPF_71546** | Transcriptional regulator MraZ |
| **KCPMINPF_71581** | CdaA regulatory protein CdaR |
| **KCPMINPF_71694** | RNA polymerase-binding transcription factor DksA |
| **KCPMINPF_71710** | Redox-sensing transcriptional repressor Rex |
| **KCPMINPF_71754** | Regulatory protein AfsR |

| | |
|---|---|
| **KCPMINPF_71856** | Oxygen regulatory protein NreC |
| **KCPMINPF_71946** | Transcriptional regulatory protein DegU |
| **KCPMINPF_71970** | Bifunctional transcriptional activator/DNA repair enzyme Ada |
| **KCPMINPF_72018** | Alkaline phosphatase synthesis transcriptional regulatory protein PhoP |
| **KCPMINPF_72243** | Transcriptional regulatory protein CpxR |
| **KCPMINPF_72391** | Transcriptional regulatory protein DegU |
| **KCPMINPF_72405** | Heat-inducible transcription repressor HrcA |
| **KCPMINPF_72596** | Transcriptional regulatory protein TcrA |
| **KCPMINPF_72614** | Zinc-specific metallo-regulatory protein |
| **KCPMINPF_72746** | N-acetylglucosamine repressor |
| **KCPMINPF_72829** | Transcriptional regulatory protein LiaR |
| **KCPMINPF_72871** | Redox-sensing transcriptional repressor Rex |
| **KCPMINPF_72888** | Transcriptional regulatory protein DegU |
| **KCPMINPF_72909** | Transcriptional regulatory protein TcrA |
| **KCPMINPF_73033** | Transcriptional activator protein CopR |
| **KCPMINPF_73136** | RNA polymerase-binding transcription factor DksA |
| **KCPMINPF_73429** | Arsenic resistance transcriptional regulator ArsR1 |
| **KCPMINPF_73591** | Bifunctional ligase/repressor BirA |
| **KCPMINPF_73671** | Transcriptional regulator MraZ |
| **KCPMINPF_73692** | Transcriptional repressor NrdR |
| **KCPMINPF_73801** | Transcriptional regulatory protein DegU |
| **KCPMINPF_73862** | Acetoin catabolism regulatory protein |
| **KCPMINPF_73939** | Ribose operon repressor |
| **KCPMINPF_73979** | Transcriptional regulatory protein OmpR |
| **KCPMINPF_74023** | Transcriptional regulatory protein TdiR |
| **KCPMINPF_74133** | Redox-sensing transcriptional repressor Rex 1 |
| **KCPMINPF_74154** | Transcriptional regulatory protein DegU |
| **KCPMINPF_74155** | Transcriptional regulatory protein LnrK |
| **KCPMINPF_74190** | LexA repressor |
| **KCPMINPF_74216** | Transcriptional regulatory protein DegU |
| **KCPMINPF_74234** | Transcriptional regulator SdrP |
| **KCPMINPF_74242** | RNA polymerase-binding transcription factor CarD |
| **KCPMINPF_74286** | DnaA regulatory inactivator Hda |
| **KCPMINPF_74312** | Transcriptional regulatory protein DegU |
| **KCPMINPF_74348** | Transcriptional regulatory protein WalR |
| **KCPMINPF_74440** | Regulatory protein AtoC |
| **KCPMINPF_74559** | Heat-inducible transcription repressor HrcA |
| **KCPMINPF_74600** | Hydrogen peroxide-inducible genes activator |
| **KCPMINPF_74674** | Transcriptional repressor NrdR |
| **KCPMINPF_74805** | Transcriptional regulatory protein AfsQ1 |
| **KCPMINPF_74827** | Bifunctional ligase/repressor BirA |
| **KCPMINPF_74829** | DNA-binding transcriptional regulator NtrC |
| **KCPMINPF_74863** | DNA-binding transcriptional activator DevR/DosR |

| | |
|---|---|
| **KCPMINPF_74986** | Regulatory protein AtoC |
| **KCPMINPF_75097** | Diphtheria toxin repressor |
| **KCPMINPF_75233** | Transcriptional regulator MraZ |
| **KCPMINPF_75347** | Transcriptional regulatory protein WalR |
| **KCPMINPF_75529** | Transcriptional regulatory protein DegU |
| **KCPMINPF_75559** | Copper-sensing transcriptional repressor CsoR |
| **KCPMINPF_75630** | Heat-inducible transcription repressor HrcA |
| **KCPMINPF_75669** | Transcriptional regulatory protein KdpE |
| **KCPMINPF_75693** | Bifunctional transcriptional activator/DNA repair enzyme Ada |
| **KCPMINPF_75783** | Transcriptional regulatory protein LiaR |
| **KCPMINPF_75806** | Transcriptional regulatory protein QseB |
| **KCPMINPF_75842** | Transcriptional regulatory protein WalR |
| **KCPMINPF_75992** | Nitrogen regulatory protein P-II 2 |
| **KCPMINPF_76032** | Transcriptional regulatory protein QseF |
| **KCPMINPF_76367** | Alkaline phosphatase synthesis transcriptional regulatory protein PhoP |
| **KCPMINPF_76377** | Alkaline phosphatase synthesis transcriptional regulatory protein SphR |
| **KCPMINPF_76405** | Transcriptional regulator MraZ |
| **KCPMINPF_76476** | LexA repressor |
| **KCPMINPF_76531** | KDP operon transcriptional regulatory protein KdpE |
| **KCPMINPF_76624** | Transcriptional regulatory protein PmpR |
| **KCPMINPF_76745** | Bifunctional ligase/repressor BirA |
| **KCPMINPF_76784** | Hydrogenase transcriptional regulatory protein hupR1 |
| **KCPMINPF_76936** | Transcriptional regulatory protein LiaR |
| **KCPMINPF_77006** | Transcriptional regulatory protein WalR |
| **KCPMINPF_77152** | LexA repressor |
| **KCPMINPF_77166** | Nitrogen regulatory protein P-II 2 |
| **KCPMINPF_77289** | flagellum biosynthesis repressor protein FlbT |
| **KCPMINPF_77343** | ATP phosphoribosyltransferase regulatory subunit |
| **KCPMINPF_77813** | Transcriptional regulatory protein QseB |
| **KCPMINPF_77936** | Nitrogen regulatory protein |
| **KCPMINPF_78096** | Iron-dependent repressor IdeR |
| **KCPMINPF_78221** | Alkaline phosphatase synthesis transcriptional regulatory protein PhoP |
| **KCPMINPF_78253** | Alkaline phosphatase synthesis transcriptional regulatory protein PhoP |
| **KCPMINPF_78290** | Redox-sensing transcriptional repressor Rex 1 |
| **KCPMINPF_78578** | Regulatory protein AtoC |
| **KCPMINPF_78614** | Transcriptional regulatory protein DegU |
| **KCPMINPF_78721** | Transcriptional regulatory protein DegU |
| **KCPMINPF_78739** | Leucine-responsive regulatory protein |
| **KCPMINPF_78793** | Transcriptional regulatory protein KdpE |
| **KCPMINPF_78883** | Transcriptional regulatory protein KdpE |
| **KCPMINPF_79061** | N-acetylglucosamine repressor |
| **KCPMINPF_79069** | Transcriptional regulatory protein DegU |
| **KCPMINPF_79184** | Transcriptional regulatory protein LiaR |

| | |
|---|---|
| **KCPMINPF_79333** | Oxygen regulatory protein NreC |
| **KCPMINPF_79378** | Bifunctional ligase/repressor BirA |
| **KCPMINPF_79506** | N-acetylglucosamine repressor |
| **KCPMINPF_79511** | N-acetylglucosamine repressor |
| **KCPMINPF_79703** | Luminescence regulatory protein LuxO |
| **KCPMINPF_79799** | Oxygen regulatory protein NreC |
| **KCPMINPF_79809** | Ribose operon repressor |
| **KCPMINPF_79975** | Regulatory protein RecX |
| **KCPMINPF_80045** | Transcriptional regulatory protein LiaR |
| **KCPMINPF_80130** | ATP phosphoribosyltransferase regulatory subunit |
| **KCPMINPF_80155** | Glycine cleavage system transcriptional activator |
| **KCPMINPF_80321** | LexA repressor |
| **KCPMINPF_80547** | Transcriptional regulatory protein LiaR |
| **KCPMINPF_80628** | Alkaline phosphatase synthesis transcriptional regulatory protein PhoP |
| **KCPMINPF_80669** | Phosphate regulon transcriptional regulatory protein PhoB |
| **KCPMINPF_80686** | Transcriptional regulatory protein WalR |
| **KCPMINPF_80922** | Transcriptional regulatory protein DegU |
| **KCPMINPF_80968** | cAMP-activated global transcriptional regulator CRP |
| **KCPMINPF_81053** | Transcriptional regulatory protein DesR |
| **KCPMINPF_81072** | RNA polymerase-binding transcription factor CarD |
| **KCPMINPF_81073** | Transcriptional regulatory protein DesR |
| **KCPMINPF_81119** | Transcriptional regulatory protein KdpE |
| **KCPMINPF_81131** | Regulatory protein AtoC |
| **KCPMINPF_81149** | RNA polymerase-binding transcription factor DksA |
| **KCPMINPF_81169** | Glucitol operon repressor |
| **KCPMINPF_81250** | Regulatory protein RecX |
| **KCPMINPF_81254** | Nitrogen regulatory protein |
| **KCPMINPF_81373** | Mercuric resistance operon regulatory protein |
| **KCPMINPF_81522** | Psp operon transcriptional activator |
| **KCPMINPF_81560** | Alkaline phosphatase synthesis transcriptional regulatory protein PhoP |
| **KCPMINPF_81561** | DNA-binding transcriptional activator DevR/DosR |
| **KCPMINPF_81781** | Transcriptional activator protein CopR |
| **KCPMINPF_81845** | Protein-arginine kinase activator protein |
| **KCPMINPF_81884** | Lactose operon repressor |
| **KCPMINPF_82008** | Heat-inducible transcription repressor HrcA |
| **KCPMINPF_82052** | Oxygen regulatory protein NreC |
| **KCPMINPF_82117** | Transcriptional regulatory protein LiaR |
| **KCPMINPF_82136** | Lactose operon repressor |
| **KCPMINPF_82176** | Transcriptional regulatory protein DegU |
| **KCPMINPF_82206** | Bifunctional ligase/repressor BirA |
| **KCPMINPF_82291** | Hydrogen peroxide-inducible genes activator |
| **KCPMINPF_82305** | Transcriptional regulatory protein DegU |
| **KCPMINPF_82551** | Glc operon transcriptional activator |

| | |
|---|---|
| **KCPMINPF_82718** | Transcriptional regulator SlyA |
| **KCPMINPF_82734** | Bifunctional transcriptional activator/DNA repair enzyme Ada |
| **KCPMINPF_82808** | Transcriptional regulatory protein ZraR |
| **KCPMINPF_82960** | PCP degradation transcriptional activation protein |
| **KCPMINPF_82993** | N-acetylglucosamine repressor |
| **KCPMINPF_83038** | Oxygen regulatory protein NreC |
| **KCPMINPF_83058** | Regulatory protein AtoC |
| **KCPMINPF_83093** | RNA polymerase-binding transcription factor CarD |
| **KCPMINPF_83238** | RNA polymerase-binding transcription factor CarD |
| **KCPMINPF_83360** | Transcriptional repressor NrdR |
| **KCPMINPF_83376** | Transcriptional regulator LsrR |
| **KCPMINPF_83467** | Transcriptional regulatory protein LnrK |
| **KCPMINPF_83504** | Oxygen regulatory protein NreC |
| **KCPMINPF_83603** | Glucitol operon repressor |
| **KCPMINPF_83605** | Transcriptional repressor IclR |
| **KCPMINPF_83633** | Oxygen regulatory protein NreC |
| **KCPMINPF_83688** | Ribose operon repressor |
| **KCPMINPF_83758** | Negative regulatory protein YxlE |
| **KCPMINPF_83772** | Oxygen regulatory protein NreC |
| **KCPMINPF_83780** | Purine catabolism regulatory protein |
| **KCPMINPF_83803** | Transcriptional regulatory protein AfsQ1 |
| **KCPMINPF_84033** | Acetoin catabolism regulatory protein |
| **KCPMINPF_84095** | Methanol dehydrogenase activator |
| **KCPMINPF_84097** | Transcriptional regulatory protein DegU |
| **KCPMINPF_84159** | Transcriptional regulatory protein LiaR |
| **KCPMINPF_84367** | Bifunctional transcriptional activator/DNA repair enzyme Ada |
| **KCPMINPF_84406** | Transcriptional repressor NrdR |
| **KCPMINPF_84412** | Murein hydrolase activator EnvC |
| **KCPMINPF_84683** | Glycerol-3-phosphate regulon repressor |
| **KCPMINPF_84723** | Heat-inducible transcription repressor HrcA |
| **KCPMINPF_84759** | Transcriptional regulatory protein WalR |
| **KCPMINPF_84816** | Transcriptional regulator |
| **KCPMINPF_84819** | Alginate biosynthesis transcriptional regulatory protein AlgB |
| **KCPMINPF_84888** | LexA repressor |
| **KCPMINPF_84889** | Transcriptional regulator LdrP |
| **KCPMINPF_85012** | Transcriptional activator protein CzcR |
| **KCPMINPF_85169** | Transcriptional regulatory protein DegU |
| **KCPMINPF_85179** | Transcriptional regulatory protein WalR |
| **KCPMINPF_85265** | Transcriptional regulatory protein WalR |
| **KCPMINPF_85344** | Oxygen regulatory protein NreC |
| **KCPMINPF_85619** | LexA repressor |
| **KCPMINPF_85628** | C4-dicarboxylate transport transcriptional regulatory protein DctD |
| **KCPMINPF_85695** | KDP operon transcriptional regulatory protein KdpE |

| | |
|---|---|
| **KCPMINPF_85737** | Pectin degradation repressor protein KdgR |
| **KCPMINPF_85851** | KDP operon transcriptional regulatory protein KdpE |
| **KCPMINPF_85894** | Transcriptional regulatory protein LiaR |
| **KCPMINPF_85914** | Regulatory protein AtoC |
| **KCPMINPF_85999** | LexA repressor |
| **KCPMINPF_86288** | N-acetylglucosamine repressor |
| **KCPMINPF_86347** | Iron-dependent repressor IdeR |
| **KCPMINPF_86371** | Bifunctional transcriptional activator/DNA repair enzyme Ada |
| **KCPMINPF_86422** | Regulatory protein AtoC |
| **KCPMINPF_86423** | Transcriptional regulatory protein DegU |
| **KCPMINPF_86900** | Bifunctional ligase/repressor BirA |
| **KCPMINPF_86924** | Regulatory protein AtoC |
| **KCPMINPF_86993** | Transcriptional regulatory protein LiaR |
| **KCPMINPF_87007** | KDP operon transcriptional regulatory protein KdpE |
| **KCPMINPF_87109** | DNA-binding transcriptional activator DecR |
| **KCPMINPF_87209** | Regulatory protein AtoC |
| **KCPMINPF_87218** | Iron-dependent repressor IdeR |
| **KCPMINPF_87342** | Regulatory protein RecX |
| **KCPMINPF_87344** | Transcriptional regulator MraZ |
| **KCPMINPF_87390** | Transcriptional regulatory protein WalR |
| **KCPMINPF_87497** | Leucine-responsive regulatory protein |
| **KCPMINPF_87679** | Transcriptional regulator PerR |
| **KCPMINPF_87762** | Transcriptional regulator PerR |
| **KCPMINPF_87810** | N-acetylglucosamine repressor |
| **KCPMINPF_87872** | Transcriptional regulatory protein LiaR |
| **KCPMINPF_87987** | Heat-inducible transcription repressor HrcA |
| **KCPMINPF_88056** | Oxygen regulatory protein NreC |
| **KCPMINPF_88072** | Transcriptional regulatory protein LiaR |
| **KCPMINPF_88075** | Purine catabolism regulatory protein |
| **KCPMINPF_88083** | Zinc-specific metallo-regulatory protein |
| **KCPMINPF_88127** | Oxygen regulatory protein NreC |
| **KCPMINPF_88140** | RNA polymerase-binding transcription factor DksA |
| **KCPMINPF_88161** | Alkaline phosphatase synthesis transcriptional regulatory protein PhoP |
| **KCPMINPF_88313** | Transcriptional regulatory protein DegU |
| **KCPMINPF_88390** | Transcriptional regulatory protein LiaR |
| **KCPMINPF_88826** | Transcriptional repressor NrdR |
| **KCPMINPF_88888** | Alkaline phosphatase synthesis transcriptional regulatory protein PhoP |
| **KCPMINPF_89084** | Transcriptional regulatory protein WalR |
| **KCPMINPF_89205** | Iron-dependent repressor IdeR |
| **KCPMINPF_89298** | Regulatory protein AtoC |
| **KCPMINPF_89558** | Heat-inducible transcription repressor HrcA |
| **KCPMINPF_89813** | Transcriptional regulatory protein DegU |
| **KCPMINPF_89821** | Alkaline phosphatase synthesis transcriptional regulatory protein PhoP |

| | |
|---|---|
| **KCPMINPF_89849** | Oxygen regulatory protein NreC |
| **KCPMINPF_90092** | Transcriptional regulatory protein QseB |
| **KCPMINPF_90179** | Transcriptional regulatory protein FixJ |
| **KCPMINPF_90243** | Transcriptional activator protein CzcR |
| **KCPMINPF_90307** | Bifunctional transcriptional activator/DNA repair enzyme Ada |
| **KCPMINPF_90319** | Transcriptional regulatory protein WalR |
| **KCPMINPF_90368** | Transcriptional regulatory protein WalR |
| **KCPMINPF_90371** | Acetoin catabolism regulatory protein |
| **KCPMINPF_90591** | Oxygen regulatory protein NreC |
| **KCPMINPF_90628** | Transcriptional regulatory protein KdpE |
| **KCPMINPF_90676** | Transcriptional regulator LsrR |
| **KCPMINPF_90699** | Transcriptional regulator AcuR |
| **KCPMINPF_90753** | Transcriptional regulatory protein BaeR |
| **KCPMINPF_90869** | KDP operon transcriptional regulatory protein KdpE |
| **KCPMINPF_90920** | Transcriptional regulatory protein QseF |
| **KCPMINPF_90955** | Transcriptional regulator PerR |
| **KCPMINPF_90984** | Bifunctional ligase/repressor BirA |
| **KCPMINPF_91187** | Photosynthetic apparatus regulatory protein RegA |
| **KCPMINPF_91188** | Trans-acting regulatory protein HvrA |
| **KCPMINPF_91199** | Transcriptional regulatory protein LiaR |
| **KCPMINPF_91347** | Oxygen regulatory protein NreC |
| **KCPMINPF_91438** | Transcriptional activator protein CopR |
| **KCPMINPF_91487** | Regulatory protein AtoC |
| **KCPMINPF_91558** | Transcriptional regulatory protein DegU |
| **KCPMINPF_91653** | Regulatory protein AtoC |
| **KCPMINPF_91661** | Transcriptional regulatory protein KdpE |
| **KCPMINPF_91669** | PTS-dependent dihydroxyacetone kinase operon regulatory protein |
| **KCPMINPF_91756** | Flagellar transcriptional regulator FlhD |
| **KCPMINPF_91757** | Flagellar transcriptional regulator FlhC |
| **KCPMINPF_91762** | Transcriptional regulatory protein ZraR |
| **KCPMINPF_91793** | Photosynthetic apparatus regulatory protein RegA |
| **KCPMINPF_91811** | Transcriptional regulatory protein LiaR |
| **KCPMINPF_92189** | Transcriptional regulatory protein AfsQ1 |
| **KCPMINPF_92281** | Transcriptional regulatory protein SrrA |
| **KCPMINPF_92433** | Glucitol operon repressor |
| **KCPMINPF_92516** | Hydrogenase transcriptional regulatory protein hupR1 |
| **KCPMINPF_92598** | Transcriptional regulatory protein LiaR |
| **KCPMINPF_92708** | Glycine cleavage system transcriptional activator |
| **KCPMINPF_92720** | Transcriptional regulator MraZ |
| **KCPMINPF_92828** | Transcriptional repressor NrdR |
| **KCPMINPF_92850** | Transcriptional regulatory protein WalR |
| **KCPMINPF_92928** | Oxygen regulatory protein NreC |
| **KCPMINPF_92934** | Transcriptional activator protein CzcR |

| | |
|---|---|
| **KCPMINPF_92949** | Transcriptional regulatory protein WalR |
| **KCPMINPF_93293** | Transcriptional regulatory protein DegU |
| **KCPMINPF_93331** | Bifunctional transcriptional activator/DNA repair enzyme Ada |
| **KCPMINPF_93402** | Transcriptional regulatory protein WalR |
| **KCPMINPF_93601** | N-acetylglucosamine repressor |
| **KCPMINPF_93673** | Oxygen regulatory protein NreC |
| **KCPMINPF_93719** | Regulatory protein AtoC |
| **KCPMINPF_93960** | Transcriptional regulatory protein LnrK |
| **KCPMINPF_94036** | Transcriptional regulatory protein DegU |
| **KCPMINPF_94038** | Oxygen regulatory protein NreC |
| **KCPMINPF_94345** | Transcriptional regulator MraZ |
| **KCPMINPF_94808** | Phosphate regulon transcriptional regulatory protein PhoB |
| **KCPMINPF_94863** | Ribose operon repressor |
| **KCPMINPF_94987** | Transcriptional regulator KdgR |
| **KCPMINPF_95120** | Transcriptional repressor SmtB |
| **KCPMINPF_95214** | Transcriptional regulatory protein RcsB |
| **KCPMINPF_95233** | Transcriptional regulatory protein LiaR |
| **KCPMINPF_95384** | Luminescence regulatory protein LuxO |
| **KCPMINPF_95761** | Redox-sensing transcriptional repressor Rex |
| **KCPMINPF_95773** | Erythritol catabolism regulatory protein EryD |
| **KCPMINPF_95856** | Transcriptional regulatory protein TcrA |
| **KCPMINPF_95869** | Alkaline phosphatase synthesis transcriptional regulatory protein SphR |
| **KCPMINPF_95889** | Alkaline phosphatase synthesis transcriptional regulatory protein PhoP |
| **KCPMINPF_95913** | Phosphate regulon transcriptional regulatory protein PhoB |
| **KCPMINPF_96173** | Transcriptional regulatory protein WalR |
| **KCPMINPF_96312** | Transcriptional regulatory protein DegU |
| **KCPMINPF_96363** | Copper-sensing transcriptional repressor CsoR |
| **KCPMINPF_96483** | Oxygen regulatory protein NreC |
| **KCPMINPF_96485** | Redox-sensing transcriptional repressor Rex 1 |
| **KCPMINPF_96507** | Oxygen regulatory protein NreC |
| **KCPMINPF_96531** | Glucitol operon repressor |
| **KCPMINPF_96631** | DNA-binding transcriptional regulator NtrC |
| **KCPMINPF_96633** | DNA-binding transcriptional regulator NtrC |
| **KCPMINPF_97044** | Transcriptional regulatory protein WalR |
| **KCPMINPF_97111** | Iron-dependent repressor IdeR |
| **KCPMINPF_97427** | Alkaline phosphatase synthesis transcriptional regulatory protein PhoP |
| **KCPMINPF_97479** | DNA-binding transcriptional activator DevR/DosR |
| **KCPMINPF_97607** | Ribose operon repressor |
| **KCPMINPF_97694** | Alkaline phosphatase synthesis transcriptional regulatory protein PhoP |
| **KCPMINPF_97699** | Transcriptional regulatory protein WalR |
| **KCPMINPF_97709** | Ribose operon repressor |
| **KCPMINPF_97739** | KDP operon transcriptional regulatory protein KdpE |
| **KCPMINPF_97743** | Transcriptional regulatory protein DegU |

| | |
|---|---|
| **KCPMINPF_97774** | Transcriptional activator protein CzcR |
| **KCPMINPF_97783** | PTS-dependent dihydroxyacetone kinase operon regulatory protein |
| **KCPMINPF_97855** | Alkaline phosphatase synthesis transcriptional regulatory protein PhoP |
| **KCPMINPF_97880** | Transcriptional regulator SlyA |
| **KCPMINPF_97886** | Copper-sensing transcriptional repressor CsoR |
| **KCPMINPF_98002** | Transcriptional regulatory protein KdpE |
| **KCPMINPF_98013** | DNA-binding transcriptional activator DevR/DosR |
| **KCPMINPF_98024** | N-acetylglucosamine repressor |
| **KCPMINPF_98107** | Bifunctional ligase/repressor BirA |
| **KCPMINPF_98151** | Bifunctional ligase/repressor BirA |
| **KCPMINPF_98319** | Oxygen regulatory protein NreC |
| **KCPMINPF_98341** | Redox-sensing transcriptional repressor Rex 1 |
| **KCPMINPF_98357** | Transcriptional regulatory protein LiaR |
| **KCPMINPF_98374** | DNA-binding transcriptional activator DevR/DosR |
| **KCPMINPF_98432** | Alkaline phosphatase synthesis transcriptional regulatory protein PhoP |
| **KCPMINPF_98456** | Purine catabolism regulatory protein |
| **KCPMINPF_98625** | Transcriptional regulatory protein DegU |
| **KCPMINPF_98642** | Alkaline phosphatase synthesis transcriptional regulatory protein PhoP |
| **KCPMINPF_98703** | Trp operon repressor |
| **KCPMINPF_98723** | Alkaline phosphatase synthesis transcriptional regulatory protein PhoP |
| **KCPMINPF_98935** | Transcriptional repressor IclR |
| **KCPMINPF_98943** | Transcriptional regulatory protein DegU |
| **KCPMINPF_98947** | Oxygen regulatory protein NreC |
| **KCPMINPF_99053** | Transcriptional regulatory protein LiaR |
| **KCPMINPF_99066** | Alkaline phosphatase synthesis transcriptional regulatory protein PhoP |
| **KCPMINPF_99198** | Leucine-responsive regulatory protein |
| **KCPMINPF_99214** | Transcriptional repressor NrdR |
| **KCPMINPF_99327** | Transcriptional regulator BlaI |
| **KCPMINPF_99342** | Transcriptional regulatory protein BasR |
| **KCPMINPF_99353** | Transcriptional regulatory protein DegU |
| **KCPMINPF_99450** | Oxygen regulatory protein NreC |
| **KCPMINPF_99576** | Transcriptional regulatory protein WalR |
| **KCPMINPF_99607** | RNA polymerase-binding transcription factor DksA |
| **KCPMINPF_99641** | Arabinose metabolism transcriptional repressor |
| **KCPMINPF_99962** | DnaA regulatory inactivator Hda |
| **KCPMINPF_99992** | Regulatory protein AtoC |
| **KCPMINPF_99994** | Transcriptional regulatory protein LiaR |
