## Additional_File_5 for "PredicTF: a tool to predict bacterial transcription factors in complex microbial communities"

---

**Additional file 5. Table S3:** Number of Transcription Factors (TFs) per TF family mapped to each of the 11 metatranscriptomes of reference from the same bioreactor where the metagenome (accession number PRJNA511011, NCBI) used to predict the putative TFs was collected. The different metatranscriptomes are represented by their European Nucleotide Archive accession numbers.

| | **SRR7091381** | **SRR7091385** | **SRR7091400** | **SRR7091401** | **SRR7091402** | **SRR7091406** | **SRR7523233** | **SRR7523243** | **SRR7523244** | **SRR7523245** | **SRR7523246** |
|---|---|---|---|---|---|---|---|---|---|---|---|
| | Collection dates (YYYY-MM-DD)[a] | | | | | | | | | | |
| **TF family** | 2016-08-11 | 2015-08-06 | 2016-06-21 | 2016-07-12 | 2016-08-30 | 2016-10-27 | 2015-11-19 | 2016-11-08 | 2016-02-12 | 2016-05-02 | 2016-11-17 |
| **OmpR/PhoB** | 40 | 18 | 16 | 37 | 29 | 15 | 19 | 2 | 15 | 13 | 25 |
| **LacI/GalR** | 31 | 6 | 14 | 19 | 16 | 8 | 9 | 0 | 9 | 9 | 9 |
| **NarL/FixJ** | 26 | 20 | 23 | 37 | 27 | 21 | 20 | 9 | 18 | 18 | 26 |
| **NtrC/DctD** | 17 | 5 | 14 | 21 | 9 | 12 | 8 | 8 | 7 | 12 | 10 |
| **Fur** | 12 | 5 | 7 | 13 | 7 | 11 | 0 | 3 | 3 | 7 | 10 |
| **LysR** | 7 | 3 | 8 | 12 | 3 | 3 | 5 | 3 | 7 | 4 | 8 |
| **LexA** | 7 | 3 | 3 | 8 | 5 | 4 | 4 | 0 | 2 | 2 | 5 |
| **GntR** | 5 | 2 | 1 | 4 | 2 | 3 | 1 | 1 | 2 | 1 | 1 |
| **CopY** | 5 | 2 | 3 | 6 | 3 | 1 | 3 | 3 | 3 | 4 | 4 |
| **IclR** | 4 | 3 | 4 | 5 | 4 | 3 | 4 | 0 | 4 | 3 | 3 |
| **RelB** | 2 | 0 | 1 | 1 | 0 | 0 | 1 | 0 | 1 | 1 | 1 |
| **MarR/SlyA** | 2 | 1 | 0 | 2 | 2 | 2 | 0 | 0 | 0 | 1 | 2 |
| **Lrp** | 2 | 1 | 1 | 3 | 0 | 2 | 1 | 1 | 2 | 2 | 3 |
| **DtxR/MntR** | 2 | 1 | 1 | 2 | 1 | 0 | 1 | 1 | 1 | 1 | 0 |
| **CsoR** | 2 | 2 | 2 | 2 | 4 | 3 | 2 | 1 | 1 | 2 | 3 |
| **AbiEi** | 2 | 1 | 1 | 2 | 2 | 2 | 0 | 0 | 1 | 0 | 0 |
| **TetR** | 1 | 0 | 0 | 0 | 0 | 0 | 0 | 0 | 0 | 0 | 1 |
| **Rrf2** | 1 | 0 | 1 | 3 | 0 | 2 | 0 | 2 | 1 | 0 | 3 |
| **LuxR** | 1 | 0 | 0 | 0 | 0 | 0 | 0 | 0 | 0 | 0 | 0 |
| **FIS** | 1 | 0 | 0 | 1 | 0 | 0 | 0 | 0 | 0 | 0 | 0 |
| **FNR/CRP** | 0 | 0 | 0 | 1 | 0 | 0 | 0 | 0 | 0 | 0 | 1 |
| **MerR** | 0 | 0 | 0 | 3 | 1 | 0 | 0 | 0 | 0 | 0 | 0 |
| **ArsR** | 0 | 0 | 0 | 2 | 1 | 0 | 1 | 0 | 1 | 0 | 0 |

[a] Collection dates of the different metatranscriptomes. YYYY, year. MM, month. DD, day.
