## Additional_File_6 for "PredicTF: a tool to predict bacterial transcription factors in complex microbial communities"

---

**Additional file 6: Fig. S4**

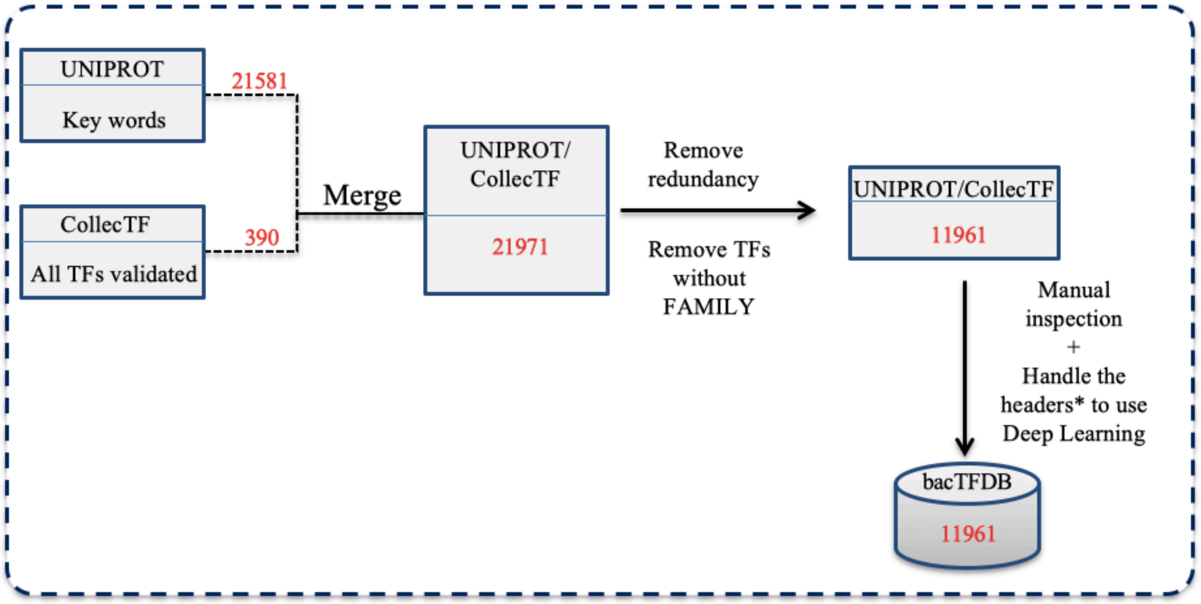


**Additional file 1: Fig. S1** Bacterial Transcription Factor Data Base (bacTFDB) were created from from two publicly available databases. We collect 390 TFs from CollecTF and 21.581 from UniProt  (accessed 8-Sep-2019) accumulating 21.581 TF amino acid

sequences. We merged the data from CollecTF and UniProt databases resulting in a total of 21.971 TFs amino acid. We removed redundant TF entries and since PredicTF was designed to also assign TF family, TF sequences lacking a TF family were removed. Finally, a manual inspection was performed to remove misleading of spelling, case sensitive and presence of characters associate to the database header. The final database (bacTFDB) contains a total of 11.691 TF unique sequences.
