## Additional_File_7 for "PredicTF: a tool to predict bacterial transcription factors in complex microbial communities"

### Additional file 7: Table S4

Description of BacTFDB subsets used to train models to predict TFs for model organisms

| Organism | Database | Description | Model |
|---|---|---|---|
| *Escherichia coli* | bacTFDB-no-coli | bacTFDB without *E. coli* transcription factors | PredicTF-no-coli |
| *Bacillus subtilis* | bacTFDB-no-subtilis | bacTFDB without *B. subtilis* transcription factors | PredicTF-no-subtilis |
| *Caulobacter crescentus* | bacTFDB-no-crescentus | bacTFDB without *C. crescentus* transcription factors | PredicTF-no-crescentus |
| *Pseudomonas fluorescens* | bacTFDB-no-fluorescens | bacTFDB without *P. fluorescens* transcription factors | PredicTF-no-fluorescens |
| *Azotobacter vinelandii* | bacTFDB-no-vinelandii | bacTFDB without *A. vinelandii* transcription factors | PredicTF-no-vinelandii |
