## Additional_File_8 for "PredicTF: a tool to predict bacterial transcription factors in complex microbial communities"

---

### Additional file 8:

Equations used to calculate PredicTF’s accuracy and performance.

*Equation 1*

$$Performance(\%) = \frac{PredictedTFs * 100}{AnnotatedTFs}$$

where, *Performance (%)* is calculated by the ratio of the total number of TFs predicted by PredicTF (*Predicted TFs*) to the total number of proteins annotated as TFs in NCBI (*Annotated TFs*) multiplied by 100.

*Equation 2*

$$Accuracy(\%) = \frac{TFspredictedcorrectly * 100}{TFspredicted}$$

where, *Accuracy (%)* is determined by the ratio of the total number of TFs predicted by PredicTF in agreement with NCBI annotation (*TFs predicted correctly*) to the total number of TFs predicted by PredicTF (*TFs predicted*) multiplied by 100.

*Equation 3*

$$Accuracy for putative TFs (\%) = \frac{putative TFs predicted correctly * 100}{putative TFs predicted}$$

where, *Accuracy for putative TFs (%)* is determined by the total number of putative TFs predicted correctly divided by putative TFs predicted multiplied by 100; *Putative TFs predicted correctly* is the total number of putative TFs predicted correctly by PredicTF in agreement with NCBI annotation; and, *Putative TFs predicted* is the total number of putative TFs predicted by PredicTF.
